# 155 years after Van Beneden: redescription and first molecular characterisation of the enigmatic type species, *Ascarophis morrhuae* Van Beneden, 1870 (Nematoda, Cystidicolidae), and comparison to other *Ascarophis* species in the North Atlantic

**DOI:** 10.64898/2026.04.15.718624

**Authors:** Ralph G. Appy, Maarten P. M. Vanhove, Ken Mackenziek, Jesús S. Hernández-Orts, Nikol Kmentová

**Author notes:** Correspondence to: Ralph G. Appy.

## Abstract

Nematodes belonging to the Cystidicolidae Skrjabin, 1946 constitute more than 23 genera of 111 recognized species in fish from many habitats including the deep-sea, continental shelves, estuarine and freshwater habitats. The taxonomy of many species within the Cystidicolidae is unsettled due to their small size and correspondingly small morphological characters requiring use of scanning electron microscopy and supported more recently by molecular studies. The type species, *Ascarophis morrhuae* Van Beneden, 1870, which belongs to one of the first described and most speciose cystidicolid genera with 46 species, is based on a two-sentence description of a single female specimen from an Atlantic cod, *Gadus morhua*, presumably captured off the coast of Belgium in the North Sea (Van Beneden, 1870). New material was collected/examined from Atlantic cod and haddock, *Melanogrammus aeglefinus*, from Iceland and the North Sea and specimens present in the Natural History Museum, London were also studied. Based on these materials, *A. morrhuae* is morphologically redescribed and the first DNA sequences of this species are provided, it is differentiated from other *Ascarophis* species present in the North Atlantic and previous records are reviewed. This information provides a foundation for taxonomic and phylogenetic reconsideration of all cystidicolid nematodes and related families.

## Introduction

The nematode family Cystidicolidae Skrjabin, 1946, which includes *Ascarophis* Van Beneden, 1870^1^ [^1^While a preponderance of authors use 1871 as the date establishing *A. morrhuae* (e.g., Nicoll, 1907; Yorke and Maplestone, 1926; Baylis, 1933; Gordon, 1951; Polyanskii, 1952; Rasheed, 1965; Moravec et al., 2024, Dollfus & Campana-Rouget (1956), Srivastava (1966), Skrjabin et al. (1967), Petter (1970) and Appy (1981) consider the date to be 1870. Here we adopt the date of 1870, which is the date appearing on the document which includes Van Beneden’s description (i.e., Van Beneden, 1870). We are unaware of any evidence that the date appearing on Van Beneden’s treatise is not the publication date, so the date of 1870 is in accordance with requirements of Article 21.2 of the International Code of Zoological Nomenclature.], comprises 27 genera (Beveridge & Moravec, 2020; Moravec et al., 2024) of poorly studied parasitic worms present worldwide in the digestive tract of fishes. They infect fishes in most aquatic environments including deep sea vents, marine and estuarine waters and fresh waters (Moravec, 2007). Where the life cycle is known, cystidicolid nematodes are transmitted by a single crustacean intermediate host or in some cases, an insect host in fresh waters (Moravec, 2007); the larvae frequently display precocity (Anderson, 2000) and have been shown to develop to adults in the intermediate host (Appy & Butterworth, 2011; Fagerholm & Butterworth, 1988). The systematics/taxonomy of cystidicolid nematodes is complex (Moravec, 2007; Moravec & Justine, 2010). In particular, genetic studies have shown that the Cystidicolidae is polyphyletic comprising genera and species intermixed with representatives of the Physalopteridae Railliet, 1893 and Acuaridae Railliet, Henry and Sisoff, 1912 which infect a variety of hosts including terrestrial mammals, fishes and birds or marine mammals (Aguilar–Aguilar et al., 2019; Černotíková et al., 2011; Choudhury & Nadler, 2018; Pereira et al., 2014; Vidal et al., 2016). Major factors contributing to the mismatch of taxonomic and genetic findings include the historic difficulty of studying the morphology of these exceedingly small worms, the paucity of species represented in genetic databases, and the ecological overlap of unrelated host groups including fish, birds and mammals, which likely became infected through host capture rather than cospeciation. In particular *Ascarophis morrhuae* Van Beneden, 1870, the type of the most speciose cystidicolid genus with 46 recognized species (Moravec et al., 2024), was only briefly described by Van Beneden (1870) from a single female worm from the stomach of the Atlantic cod, *Gadus morhua* Linnaeus, 1758, presumedly collected off the coast of Belgium.

Since no type specimens of *A. morrhuae* have ever been found/deposited, all subsequent descriptions of species belonging to *Ascarophis* are based on presumptive morphological characteristics, particularly details of the taxonomically important anterior end, and only four of the 46 species have been characterized genetically (Aguilar et al., 2019). Indeed, the establishment of subgenera of *Ascarophis* (i.e., *A*. (*Ascarophis*), *A*. (*Similascarophis*) and *A*. (*Dentiascarophis*) by Moravec & Nagasawa (2018) was done without the benefit of morphological details of the oral structure of the type species of the genus, *A. morrhuae*.

Additionally, as pointed out by Moravec & González-Solis (2007), the systematic value of the tiny oral features awaits additional studies combining the use of scanning electron microscopy (SEM) and genetic studies, especially including examination of new material of the type species, *A. morrhuae,* from the type host and type locality (Dollfus, 1953; Moravec & Nagasawa, 2018).

During a parasitological study of fish in the Northeast Atlantic, presumptive specimens of *A. morrhuae* were collected from haddock, *Melanogrammus aeglefinus* (Linnaeus, 1758) caught commercially in Icelandic waters and Atlantic cod and haddock collected in the North Sea by a research ship. The purpose of this study is to use these specimens to establish the morphological identity of *A. morrhuae* and provide the first genetic data for this species; in the absence of type material, to establish neotypes; and to compare *A. morrhuae* with other species of *Ascarophis* described from similar North Atlantic latitudes, including specimens available in museums and in the first author’s personal collection. Owing to its status as one of the first cystidicolid species and genus to be identified, being the most specious genus and its basal position in the Spiromorpha (Meldal et al., 2007), the morphological and genetic identity of *A. morrhuae* is a necessary foundation for the placement among the three subgenera established by Moravec & Nagasawa (2018) and subsequent taxonomic and phylogenetic analysis of the Cystidicolidae and related families.

## Materials and methods

### Collection, sample preparation, and conservation

Fresh fish were obtained from various sources including retail and wholesale markets in Ostend, Belgium, trawls conducted by Flanders Marine Institute RV Simon Stevin along the Belgian coast and frozen hosts collected by the Marine Scotland FRS *Scotia.* Fresh dead cod (N=2) and haddock (N=5) imported from Iceland, were obtained from the Vishandel Neptunus wholesale market (Ostend, Belgium). Living nematode parasites collected for morphological study (Table 1) were cleaned in saline, heat killed with hot water and preserved in 70% molecular grade ethanol. These specimens, preserved material from museums (see Table 1) and specimens present in the first author’s collection from various hosts in the Western Atlantic, were cleared in either glycerine alcohol or lactophenol. In addition, the tails of some living male specimens were viewed alive or placed in sodium dodecyl sulfate (SDS) (Wong et al., 2006) in order to visualize spicule detail. For the genetic part of the study, specimens or parts of specimens were preserved in 95% molecular grade ethanol. Measurements were made of adult worms which we defined as females with larvated eggs and males with sclerotized spicules. Morphological study was also carried out on specimens from the Natural History Museum, London (NHMUK), the U.S. National Museum of Natural History (Smithsonian Institution, Washington, D.C.) (USNM) and from the first author’s personal collection. Neotype and voucher specimens have been deposited in the NHMUK, USNM and Hasselt University (HU) collections. Accession numbers of specimens examined in the present study are provided in Table 1.

**Table 1.**
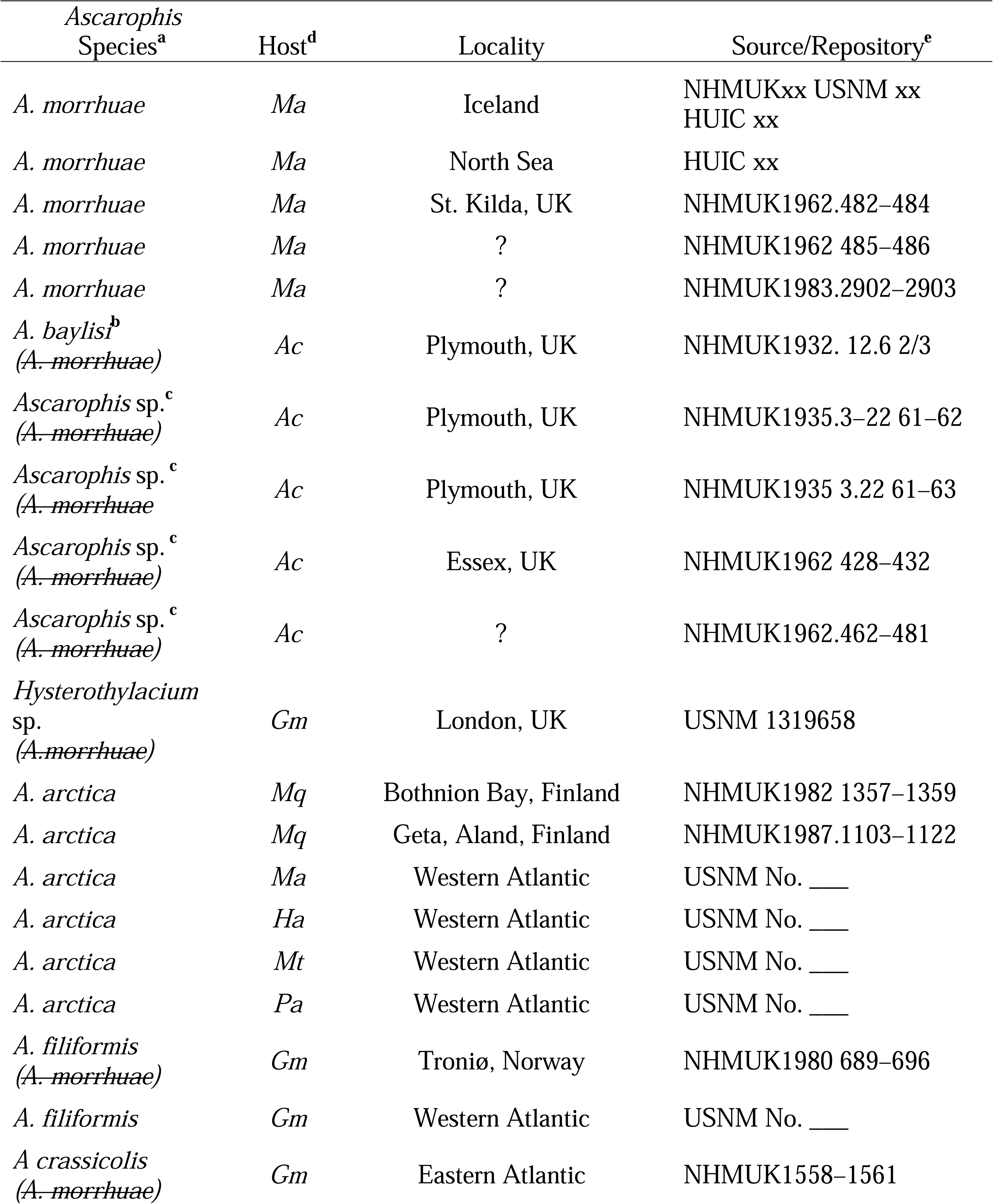
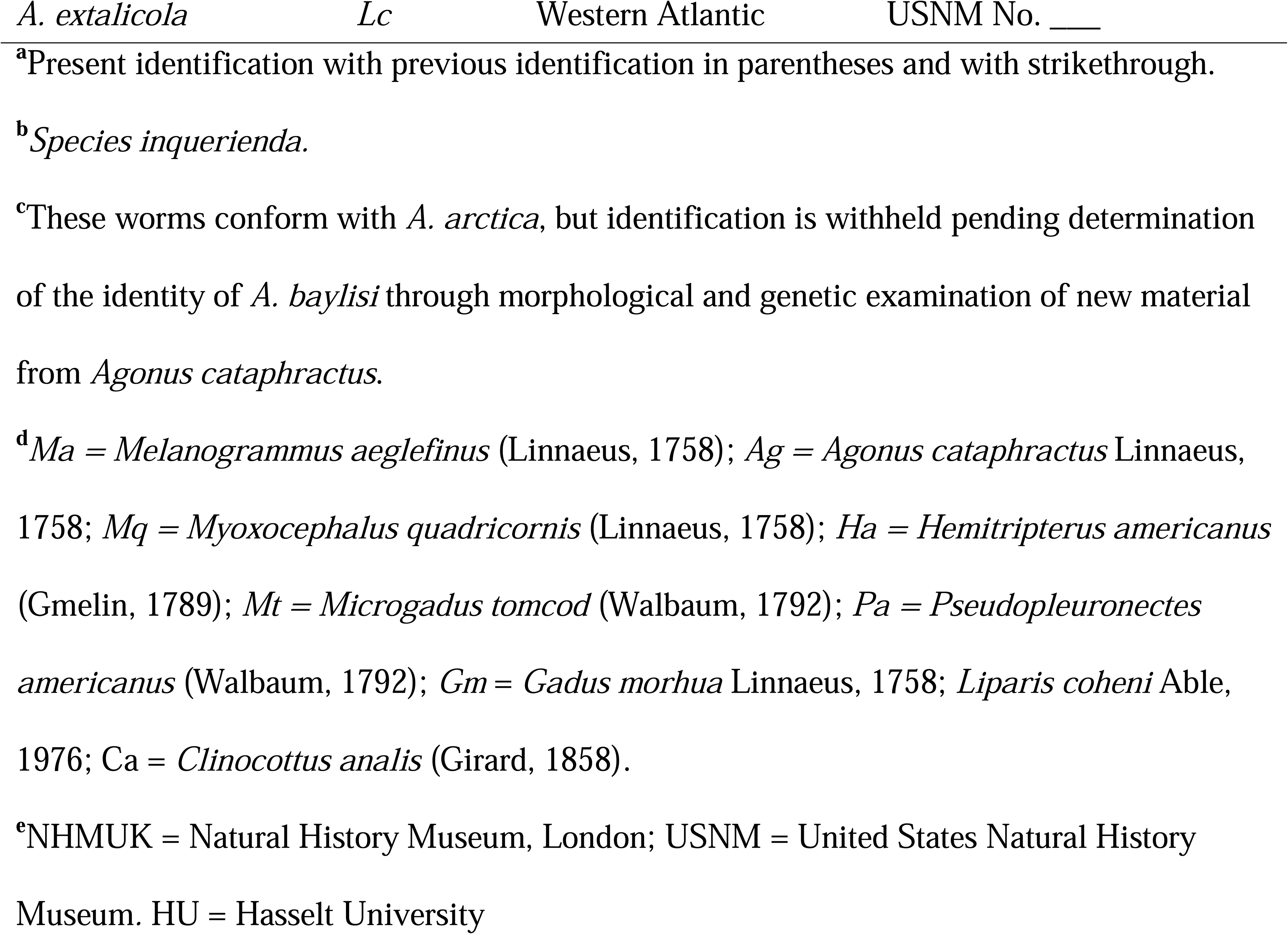
Specimens of *Ascarophis* examined in the present study including revised

### Microscopy, illustrations and statistical treatment

Visualization and photography of fresh and prepared specimens were carried out on a Leica DM2500 and a Wild M12 microscopes using bright field, phase contrast and differential interference contrast (DIC) micrography. Specimens were measured with the measure tool in Adobe Illustrator from drawings or photographs and converted from units to micrometers (μm). Drawings were made with the aid of a drawing tube, imported into and drawn/traced with Adobe Illustrator. Following morphological study, specimens were removed from slides and placed in a vial containing glycerine alcohol for long-term storage. Specimens for SEM, which were previously observed/measured in glycerine, were removed from slides, hydrated and placed in 10% formalin over–night, post fixed in 2% osmium tetroxide, and dehydrated in an ethanol series and dried with CO_2_ in a Tousimis Samdri–PVT–3D critical point drier and placed on stubs covered with adhesive copper tape. Specimens were coated with gold/palladium using a Pelco SC–4 4000 sputter coater and imaged using a Phenom Pro’X G6 Desktop SEM. The brightness, contrast and background of some electron micrographs were enhanced using Photoshop. Nematode terminology follows Anderson et al. (2009). Numerical and statistical analyses, including Analysis of Variance and Games-Howell post-hoc multiple comparisons test, were carried out using XLSTAT (Version 2023.3.1) on an iMac Studio.

### Molecular characterization and genetic distances

Whole genomic DNA extraction of three specimens of *A. morrhuae* was performed following a protocol presented in Kmentová et al. (2021). A portion of 18S rDNA region was amplified using forward primer 1073 (5–CGGGGGGAGTATGGTTGC–3) and 18SR_Saiki (5–TGATCCTTCTGCAGGTTCACCTAC–3) (Saiki et al., 1988) under the following conditions: initial denaturation at 94 °C for 4 min, then 25 cycles at 94 °C for 30 sec, followed by primer annealing at 60 °C for 30 sec, 72 °C for 1 min 30 sec and final elongation at 72 °C for 5 min. A portion of 28S rDNA region was amplified using forward primer Nem28SF (5–AGCGGAGGAAAAGAAACTAA–3) and reverse primer Nem28SR (5–TCGGAAGGAACCAGCTACTA–3) (Nadler et al., 2000) under the following conditions: initial denaturation at 94 °C for 3 min, then 33 cycles at 94 °C for 30 sec, followed by primer annealing at 55 °C for 20 sec, 72 °C for 45 sec and final elongation at 72 °C for 7 min. The reaction mix in both reactions contained one unit Mango Mix 2x Polymerase Buffer (Meridian Bioscience) containing 0.2 mM of dNTPs, 3.5 mM MgCl_2_, 0.5 mM of each primer and 2 μL of 10x diluted DNA isolate (concentration was not measured) in a total reaction volume of 25 μL. A primer cocktail for the cytochrome *c* oxidase subunit I (COX1 mtDNA) developed by Prosser et al., (2013) using the recommended PCR conditions of: initial denaturation at 94 °C for 1 min, then 5 cycles at 94 °C for 40 sec, followed by primer annealing at 45 °C for 40 sec and an elongation at 72 °C for 1 min, followed by 35 cycles starting at 94 °C for 40 sec, proceeding with primer annealing at 51 °C for 40 sec and final elongation at 72 °C for 1 min. The final elongation step consisted of 72 °C for 5 min. PCR products were gel extracted and purified using Invitrogen Gel extraction kit. Sanger sequencing was outsourced using the same primers mix as in the initial PCR reaction. The newly obtained sequences were deposited in NCBI GenBank under the accession numbers xx–xx (18S rDNA), xx–xx (28S rDNA) and xx (COX1 mtDNA).

### Phylogenetic placement

Representatives of Cystidicolidae and other families present within the cystidicolid clades based on Sokolov et al. (2019) were selected for phylogenetic reconstruction (n=48, Table 2). In addition, a portion of the 18S sequence of *Ascarophis (Similascarophis)*sp. from the wooley sculpin, *Clinocottus analis* (Girard, 1858) collected by RGA in Southern California, was included. In case of multiple sequences per species, the longest available sequence in the GenBank database was selected. Sequences were aligned in Geneious v2025.0.2 using Muscle v5.1 under the Parallel Perturbed Probcons algorithm and with four threads (Edgar, 2004) in Geneious v2025.0.2. Poorly aligned positions and divergent regions were removed with trimAl v.1.3 using the “gappy out” option. The two trimmed alignments were concatenated in Geneious v2025.0.2. The final alignments consisted of 1,568 bp (18S rDNA) and 3,174 bp (28S rDNA) including gaps and were concatenated in Geneious. The optimal substitution models was selected according to the Bayesian information criterion as implemented in ModelFinder in IQ–Tree (Kalyaanamoorthy et al., 2017) being TPM3+I+R2 and TIM3e+I+R3 for 18S rDNA and partitions, respectively. Tree topologies were estimated through Bayesian inference (BI) and maximum likelihood (ML) methods using, respectively, MrBayes v3.2.6 (Ronquist et al., 2012) on the CIPRES Science Gateway online server (Miller et al., 2010) and IQ–Tree v1.6.12 (Nguyen et al., 2015) applying the selected substitution model. Species belonging to the Physalopteridae Railliet, 1893, *Turgida torresi* Travassos, 1920 (KY990020.1, EF180069.1) and *Physaloptera sibirica* Petrow and Gorbunov, 1931 (OQ846909.1, OQ846902.1), were used as outgroup, following results of Sokolov et al. (2019).

**Table 2.** GenBank accession numbers of 18S and 28S rDNA sequences used for the phylogenetic reconstruction in the present study.

| Nematode species | GenBank Accession Number |  |
| --- | --- | --- |
|  | 28S rDNA | 18S rDNA |
| *Ascarophis adioryx* | EF079120.1 | DQ094172.1 |
| *Ascarophis morrhuae* - Iceland | XX (Present Study) | XX (Present Study) |
| *Ascarophis morrhuae* - Scotland | XX (Present Study) | XX (Present Study) |
| *Ascarophis (Similascarophis)* sp. | XX (Present Study) | XX Present Study) |
| *Ascarophis (Simmilascarphis)*<br>*morronei* |  | MK714126.1 |
| *Capillospirura ovotrichuria* |  | MN294783.1 |
| *Capillospirura* sp. |  | MG594289.1 |
| *Comephoronema oschmarini* |  | MN294782.1 |
| *Comephoronema werestschagini* |  | MN294781.1 |
| *Crassicauda boopis* |  | KY263809.1 |
| *Crassicauda magna* |  | KX354835.1 |
| *Cystidicola farionis* | MT086834.1 | MT735337.1 |
| *Cystidicola stigmatura* | AY161298.1 |  |
| *Cystidicoloides vaucheri* | KY558631.1 | KY558630.1 |
| *Echinuria borealis* |  | EF180064.1 |
| *Heliconema longissimum* |  | JF803949.1 |
| *Metabronema magnum* |  | JF803918.1 |
| *Metabronema* sp. |  | KC222020.1 |
| *Neoascarophis longispicula* |  | JF803921.1 |
| *Neoascarophis macrouri* |  | DQ442660.1 |
| *Proleptus obtusus* | KY411576.1 | KY411575.1 |
| *Rhabdochona acuminata* | MK341681.1 | MK341647.1 |
| *Rhabdochona adentata* | MN912352.1 |  |
| *Rhabdochona canadensis* | MK353494.1 | MK353466.1 |
| *Rhabdochona coronacauda* |  | PQ066505.1 |
| *Rhabdochona denudata* |  | DQ442659.1 |
| *Rhabdochona gendrei* | OR096361.1 |  |
| *Rhabdochona guerreroensis* | MG189933.1 | JF934732.1 |
| *Rhabdochona hellichi hellichi* |  | JF803913.1 |
| *Rhabdochona hospeti* | OK486360.1 | JF803938.1 |
| *Rhabdochona ictaluri* | MK353492.1 | MK353467.1 |
| *Rhabdochona kidderi kidderi* | MK353490.1 | MH754715.1 |
| *Rhabdochona kisutchi* |  | MG594298.1 |
| *Rhabdochona lichtenfelsi* | MK341682.1 | MG594300.1 |
| *Rhabdochona mazeedi* |  | JF803936.1 |
| *Rhabdochona mexicana* | MK341655.1 | MK353465.1 |
| *Rhabdochona milleri* |  | MG594294.1 |
| *Rhabdochona osorioi* | MK341676.1 | MK341648.1 |
| *Rhabdochona salgadoi* | MK341684.1 | MK341650.1 |
| *Rhabdochona xiphophori* | MK353496.1 | MK353468.1 |
| *Salmonema ephemeridarum* | MT086835.1 | JF803927.1 |
| *Spinitectus carolini* |  | DQ503464.1 |
| *Spinitectus gracilis* |  | MG594297.1 |
| *Spinitectus macrospinosus* |  | MG594296.1 |
| *Spinitectus mexicanus* | MK341687.1 | MK341639.1 |
| *Spinitectus petterae* | OP802566.1 | OP800412.1 |
| *Spinitectus tabascoensis* |  | JF803922.1 |
| *Stegophorus macronectes* | HE793715.1 | HE793715.1 |
| *Synhimantus hamatus* |  | EU004819.1 |
| *Synhimantus laticeps* |  | EU004818.1 |

In BI, two parallel runs and four chains of Metropolis–coupled Markov chain Monte Carlo iterations, run for 30 million generations with a burn-in fraction of 0.25, were employed, sampling trees every 1000th generation. Convergence was assessed by the average standard deviation of split frequencies (<0.01 in all datasets) and the effective sample size (>200) using Tracer v1.7. (Rambaut et al., 2018). Branch support values for the ML analysis were estimated using ultrafast bootstrap approximation (Hoang et al., 2018) and Shimodaira–Hasegawa–like approximate likelihood ratio tests (SH–aLRT) (Guindon et al., 2010) with 10,000 replicates for both support values (as recommended in the IQ–Tree manual). The resulting tree topologies were visualised in FigTree v1.4.4 (Rambaut, 2018) and graphically refined using Inkscape v1.4.3 (Inkscape Project. (2025). Inkscape. Retrieved from https://inkscape.org).

Genetic distances (p-distance) between all species present in the clade with *A. morrhuae* including the unpublished sequence of *A.* (*Similascarophis*) sp. collected by RGA from the wooley sculpin in Southern California (xx GenBank accession number) were calculated using MEGA for each of the studied markers separately (if available) and are presented in Table 7.

## Results

### Taxonomic account/redescription

NEMATODA DIESING, 1861

CHROMADOREA Inglis, 1983

RHABITIDA Chitwood, 1933

SPIRURINA Railliet & Henry, 1915

HABRONEMATOIDEA Ivaschkin, 1961

CYSTIDICOLIDAE Skryabin, 1946

*Ascarophis* Van Beneden 1870

Type species: *Ascarophis morrhuae* Van Beneden, 1870

Syn: *Ascaropsis morrhuae* (Van Beneden, 1870) Power and Sedgwick, 1880

NEOHOLOTYPE: NHMUK _____

NEOALLOTYPE: NHMUK _____

NEOPARATYPES (N=2): NHMUK _____ ; USNM _____; HU _____

NEOHOLOGENEPHORE _____HU

ADDITIONAL MATERIAL: See Table 1.

GENBANK: See Table 2.

NEOTYPE HOST: Haddock, *Melanogrammus aglefinus* (Linnaeus, 1758) (Gadiformes: Gadidae) (type host was *Gadus morhua* Linnaeus, 1758).

NEOTYPE LOCALITY: Iceland, Central North Atlantic (type locality was North Sea off cast of Belgium).

HABITAT: mucus of pyloric stomach.

PREVALENCE AND INTENSITY: Of five haddock examined from Iceland all were infected with one to five worms and none of two cod were infected. One of 21 haddock and two of 52 cod from the northern North Sea were infected.

ADDITIONAL HOSTS AND LOCALITIES: *Gadus morhua* Linnaeus, 1758: Barents Sea (Polyanskii, 1952); North Sea (Nicoll, 1907; Gordon, 1951); Norway (Berland, 1961); Northeast Atlantic including Iceland (Perdiguero-Alonso et al., 2008) in part, since it seems likely some of these worms may be *A. arctica,* which these authors did not report as being present in 826 cod examined. *Melanogrammus aeglefinus* (Linnaeus, 1758): North Sea (Nicoll, 1907; Gordon, 1951; Ko 1986); Barents Sea (Polyanskii, 1952). *Ciliata mustela* (Linnaeus, 1758): Aberystwyth, U.K. (Rees, 1945); Swansea, U.K. (Srivastava, 1966).

HOSTS AND LOCALITY REPORTS REQUIRING CONFIRMATION: Linstow (1900) from *Gadus morhua*, Arctic/Subarctic; Baylis (1939) from *Chelidonichthys lastoviza* (Bonnaterre, 1788) and *Agonus cataphractus* (Linnaeus, 1758), S. Devon, UK; Hanek & Threlfall (1970) from *Gasterosteus aculeatus* Linnaeus, 1758, Labrador & Newfoundland, Canada; Gaevskaya & Umnova (1977): from *Gadus morhua* Northwest Atlantic; Zubchenko (1980) from *Glyptocephalus cynoglossus* (Linnaeus, 1758), *Anarhichas lupus* Linnaeus, 1758, Northwest Atlantic; Zubchenko & Karasev (1986) from *Gadus morhua*, Barents Sea; Zubchenko (1987), specific host cannot be determined from paper; North Atlantic; Kirjušina & Vismanis (2007) from *G. morhua*, *Myoxocephalus quadricornis* (Linnaeus, 1758) and *Taurulus bubalis* (Euphrasen, 1786) from Baltic Sea off Latvia; Hogans et al. (1993) from *Alosa sapidissima* (Wilson, 1811), New Brunswick, Canada; Rodrigues et al. (1973) from *Beryx decadactylus* Cuvier, 1829, Coast of Portugal and Atlantic coast of North Africa; Cheetham & Fives (1990) from *Pholis gunnellus* (Linnaeus, 1758), Galway Bay, Ireland; Kirjušina & Vismanis (2007) from *Zoarces viviparus* (Linnaeus, 1758) and *G. morhua* from Riga area off Latvia; Koi (2012) from gadoids off Faroe Islands, Northeastern Atlantic; Sobecka & Luczak (2015) from *Zoarces viviparus* (Linnaeus, 1758) in the Baltic Sea off Oder estuary, Poland.

ERRONEOUS HOSTS AND LOCALITY REPORTS: Baylis (1933) from *Chelidonichthys lastoviza* (Bonnaterre, 1788), Plymouth, United Kingdom; Punt (1947) from *Agonus cataphractus* (Linnaeus, 1758) and *Ciliata mustela* (Linnaeus, 1758), North Sea; Petter (1970) from *Trisopterus luscus* (Linnaeus, 1758) Nantaise, France; Ko (1986) from *Agonus cataphractus* (Linnaeus, 1758), United Kingdom, Burnham-on-Crouch, United Kingdom, NHMUK 462-481; Køi et al. (2008) from *Arctogadus glacialis* (Peters, 1872) off Greenland.

DIAGNOSIS: *Ascarophis* (*Ascarophis*) with narrow pseudolabia with conical projection, and well-defined submedian labia and unindented sublabia. Deirids small, simple/unbifrucated. Body length not exceeding 5 mm in males and 8 mm in females, with overlapping transverse striations beginning abruptly at nerve ring and extending to midbody. Males with tessellated area rugosa, and ten pairs of caudal papillae; four pairs precloacal, six pairs postcloacal; papillae pedunculate except sessile tenth pair appearing. Spicules dissimilar; left spicule 500-600 μm long with thin tapered tip, right spicule 91-106 μm long. Vulva present at midbody; eggs elliptical with two broad filaments and sometimes a thin third and rarely a fourth filament emanating from a prominent knob at one pole.

**Redescription** (Tables 1–6) Figs. 1– 5)

**Figure 1.**
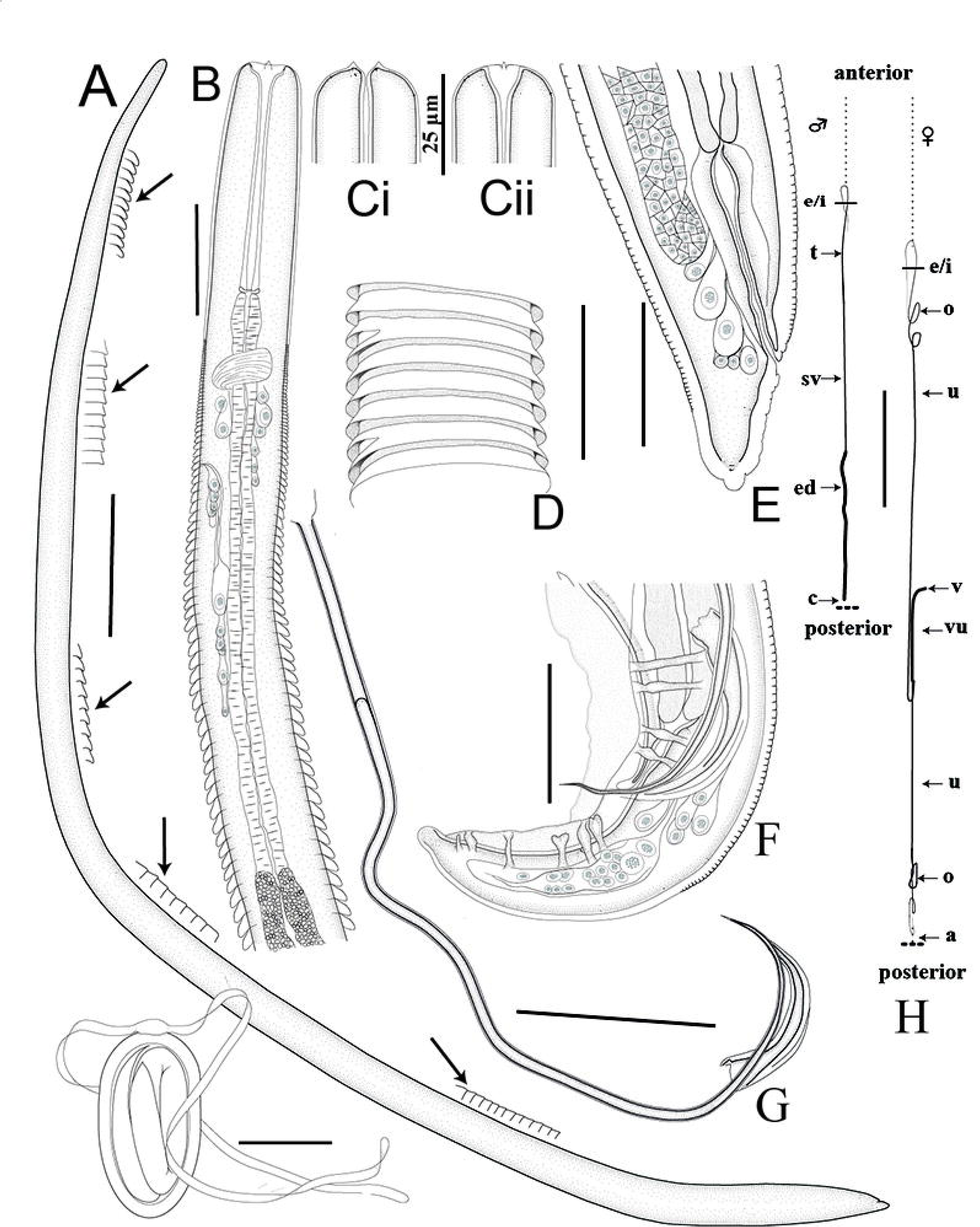
Line drawings of *Ascrophis morrhuae* Van Beneden, 1870 from the the haddock, *Melanogrammus aeglefinus* (Linnaeus, 1758) from Iceland. A. Habitus showing variation in striations along body length (see arrows) Scale bar: 0.5 mm. B. Anterior extremity, lateral view. Scale bar: 50 μm. C. Anterior end in ventral (i) and lateral (ii) views. Scale bar: 25 μm. D. Striations at the level of junction of muscular and glandular esophagus. Scale bar: 50 μm E. Posterior extremity of female, lateral view. Scale bar: 50 μm. F. Posterior extremity of male, lateral view. Scale bar: 50 μm. G. Spicules. Scale bar: 10 μm. H. Reproductive systems of male and female worms to scale. Scale bar: 1 mm. I. Mature egg. Scale Bar: 25 μm. Abbreviations: e = esophagus; ej = ejaculatory duct; i – intestine; o = ovary, sv = seminal vesicle; t = testis; u = uterus; v = vagina; vu = vagina uterina.

**Table 3.**
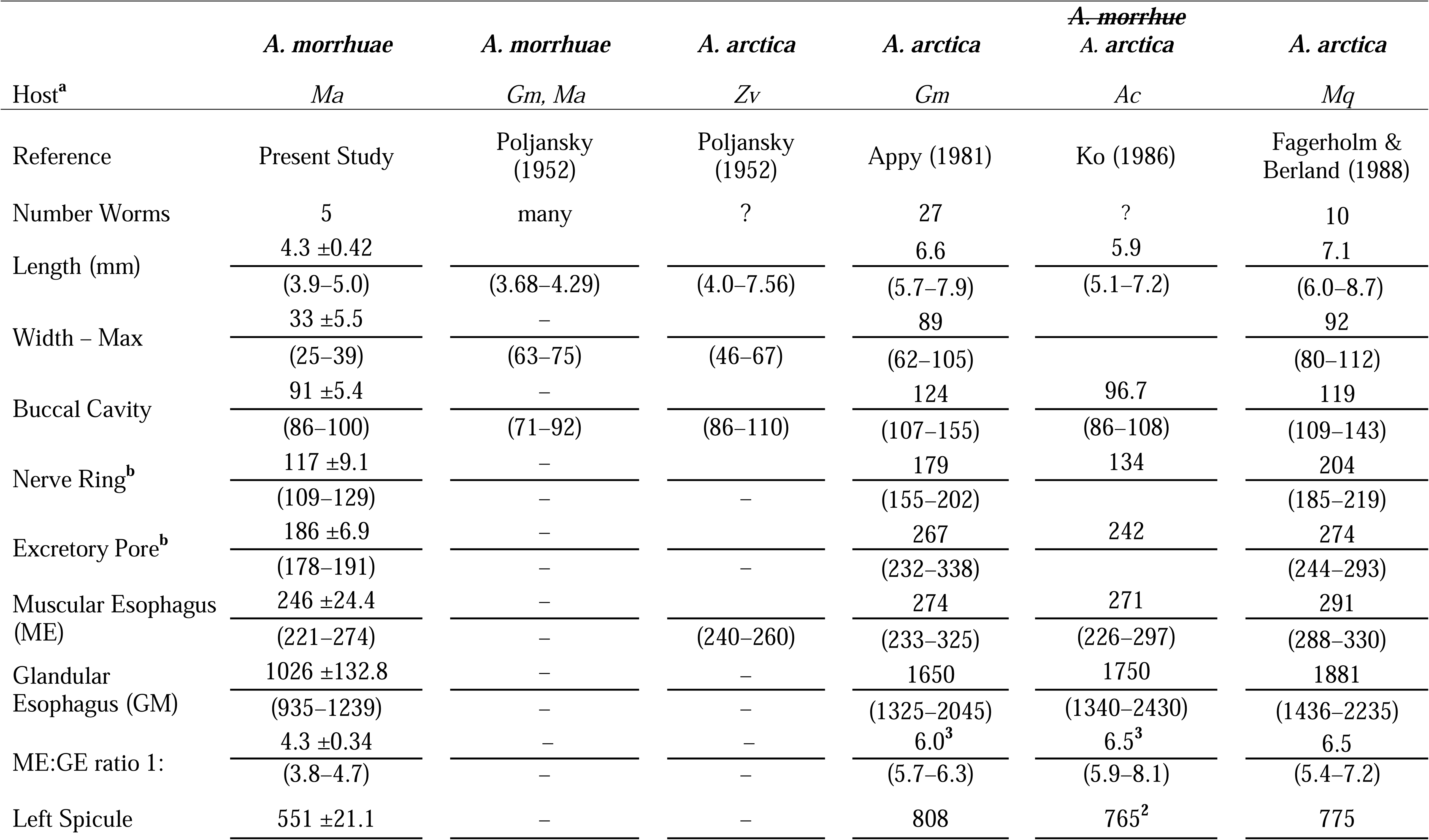
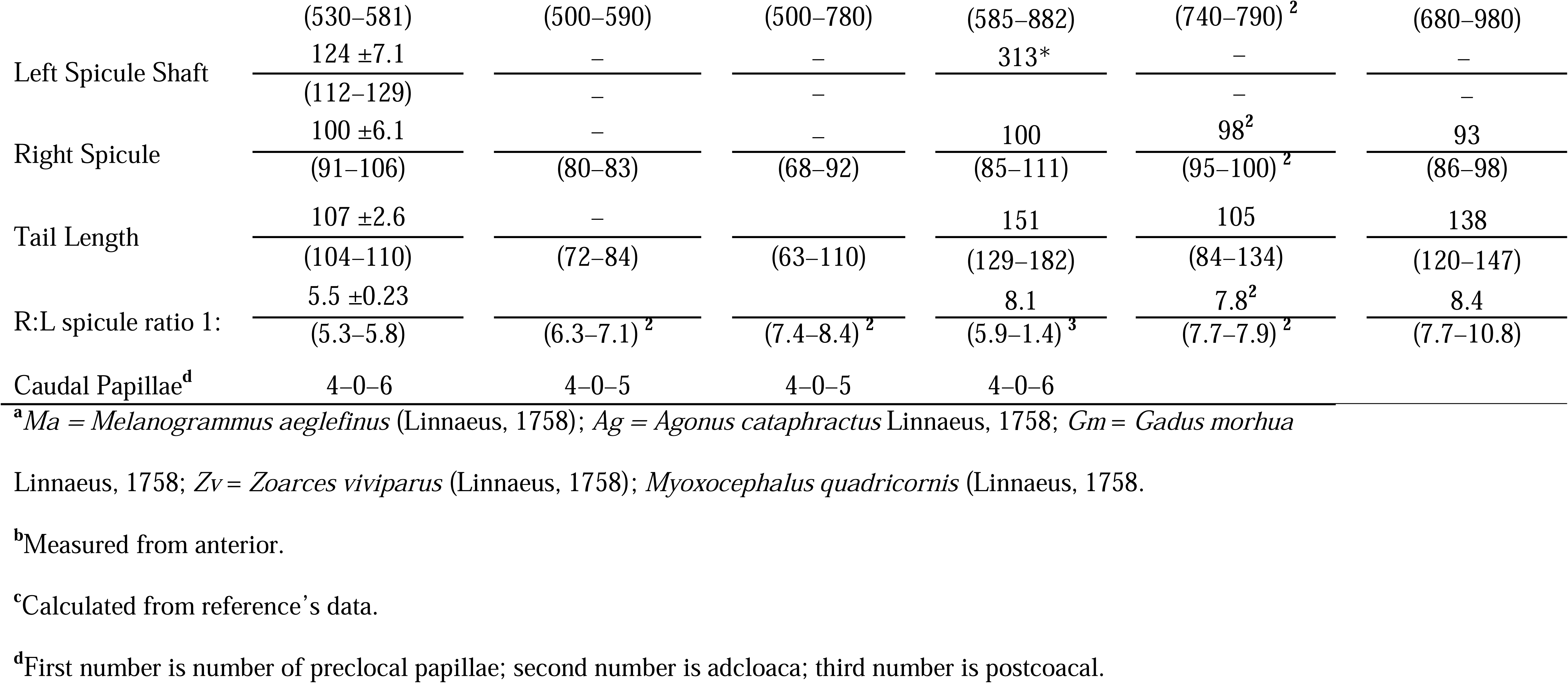
Measurements of male *Ascarophis morrhuae* Van Beneden, 1870 from the present study and *A. morrhuae* and *A. arctica* Poljansky, 1952 presented in the literature. Measurements in micrometers (μm) unless stated otherwise.

**Table 4.** Measurements of female *Ascarophis morrhuae* Van Beneden, 1870 from the present study and *A. morrhuae* and *A. arctica* Poljansky, 1952, presented in the literature. Measurements in micrometers (μm) unless stated otherwise.

| Host<sup>a</sup> | *A. morrhuae* | *A. morrhuae* | *A. arctica* | *A. arctica* | ~~*A. morrhuae*~~<br>*A. arctica* | *A. arctica* |
| --- | --- | --- | --- | --- | --- | --- |
|  | *Ma* | *Gm, Ma* | *Zv* | *Gm* | *Ac* | *Mq* |
| Reference | Present Study | Poljansky, 1952 | Poljansky, 1952 | Appy, 1981 | Ko, 1986 | Fagerholm & Berland, 1988 |
| Number of Worms | 6 | many | ? | 27 | ? | 10 |
| Length (mm) | $7.1 \pm 0.96$<br>(5.5–7.9) | –<br>(5.36–7.24) | –<br>(7.2–11.6) | 13.3<br>(10.5–15.4) | 10.8<br>(8.50–12.7) | 10.9<br>(9.7–13.0) |
| Width – Max | $127 \pm 18.4$<br>(97–147) | –<br>(93–140) | –<br>(67–114) | 124<br>(92–157) | – | 105<br>(77–142) |
| Buccal Cavity | $106 \pm 11.2$<br>(94–119) | –<br>(72–92) | –<br>(100–110) | 134<br>(110–144) | 98<br>(84–115) | 126<br>(105–138) |
| Nerve Ring<sup>b</sup> | $134 \pm 12.2$<br>(117–153) | –<br>– | –<br>– | 194<br>(172–221) | 140<br>(113–178) | 214<br>(201–233) |
| Excretory Pore<sup>b</sup> | $207 \pm 36.4$<br>(168–275) | –<br>– | –<br>– | 295<br>(269–338) | 212<br>(161–240) | 282<br>(263–335) |
| Muscular Esophagus | $297 \pm 35.8$<br>(245–344) | –<br>250 | –<br>(280–300) | 321<br>(235–414) | 246<br>(216–288) | 342<br>(307–403) |
| Glandular Esophagus | $1224 \pm 149.5$<br>(1035–1402) | –<br>(980–1470)<sup>c</sup> | –<br>(2120–3200)<sup>c</sup> | 1928<br>(1474–2554) | 1900<br>(1100–2300) | 1728<br>(1200–2295) |
| ME:GE ratio 1: | $4.1 \pm 9.45$<br>(3.4–4.8) | –<br>(3.9–5.9)<sup>c</sup> | –<br>(7.6–10.7)<sup>c</sup> | 6.0<sup>c</sup><br>(5.8–6.2) | 7.8<sup>c</sup><br>(5.0–8.0) | 5.1<br>(3.1–6.7) |
| Vulva<sup>b</sup> (mm) | $4.2 \pm 0.35$<br>(3.7–4.6) | –<br>(3.1–4.0)<sup>c</sup> | –<br>(4.0–6.0)<sup>c</sup> | 7<br>(5.5–8.3) | 5.5<br>(3.8–6.7) | 6.6<br>(5.7–7.7) |
| TL:Vulva (% body) <sup>b</sup> | $\frac{61.2 \pm 7.23}{(56.7-66.8)}$ | $\frac{-}{(55.2-55.4)^c}$ | $\frac{-}{(51.7-55.6)^c}$ | $\frac{52.6^c}{(52.3-53.9)}$ | $\frac{50.9^c}{(44.7-52.8)}$ | $\frac{60.8}{(55-64)}$ |
| Egg Length | $\frac{47 \pm 0.8}{(46-48)}$ | $\frac{-}{(41-45)}$ | $\frac{-}{(40-43)}$ | $\frac{45}{(43-48)}$ | $\frac{36}{(26-46)}$ | $\frac{47}{(45-50)}$ |
| Egg Width | $\frac{27 \pm 1.3}{(26-30)}$ | $\frac{-}{(22-27)}$ | $\frac{-}{(21-23)}$ | $\frac{24}{(22-27)}$ | $\frac{24}{(22-26)}$ | $\frac{26}{(24-29)}$ |
| Egg Filaments | 2 on 1 pole | 2 on 1 pole | many at 2 poles | many at 2 poles | Many at 2 poles<sup>d</sup> | many at 2 poles |
| Tail Length | $\frac{43 \pm 9.1}{(33-59)}$ | $\frac{-}{(34-46)}$ | $\frac{-}{(38-42)}$ | $\frac{52}{(34-71)}$ | $\frac{116}{(72-146)}$ | $\frac{44}{(35-49)}$ |

<sup>a</sup>*Ma* = *Melanogrammus aeglefinus* (Linnaeus, 1758); *Ac* = *Agonus cataphractus* Linnaeus, 1758; *Gm* = *Gadus morhua* Linnaeus, 1758; *Zv* = *Zoarcetes viviparus* (Linnaeus, 1758); *Myoxocephalus quadricornis* (Linnaeus, 1758).
<sup>b</sup>Distance from anterior end.
<sup>c</sup>Calculated from reference's data.
<sup>d</sup>Based on personal observations of material in the BMNH vs Ko's (1986) observation of 2 filaments at one pole.

**Table 5.**
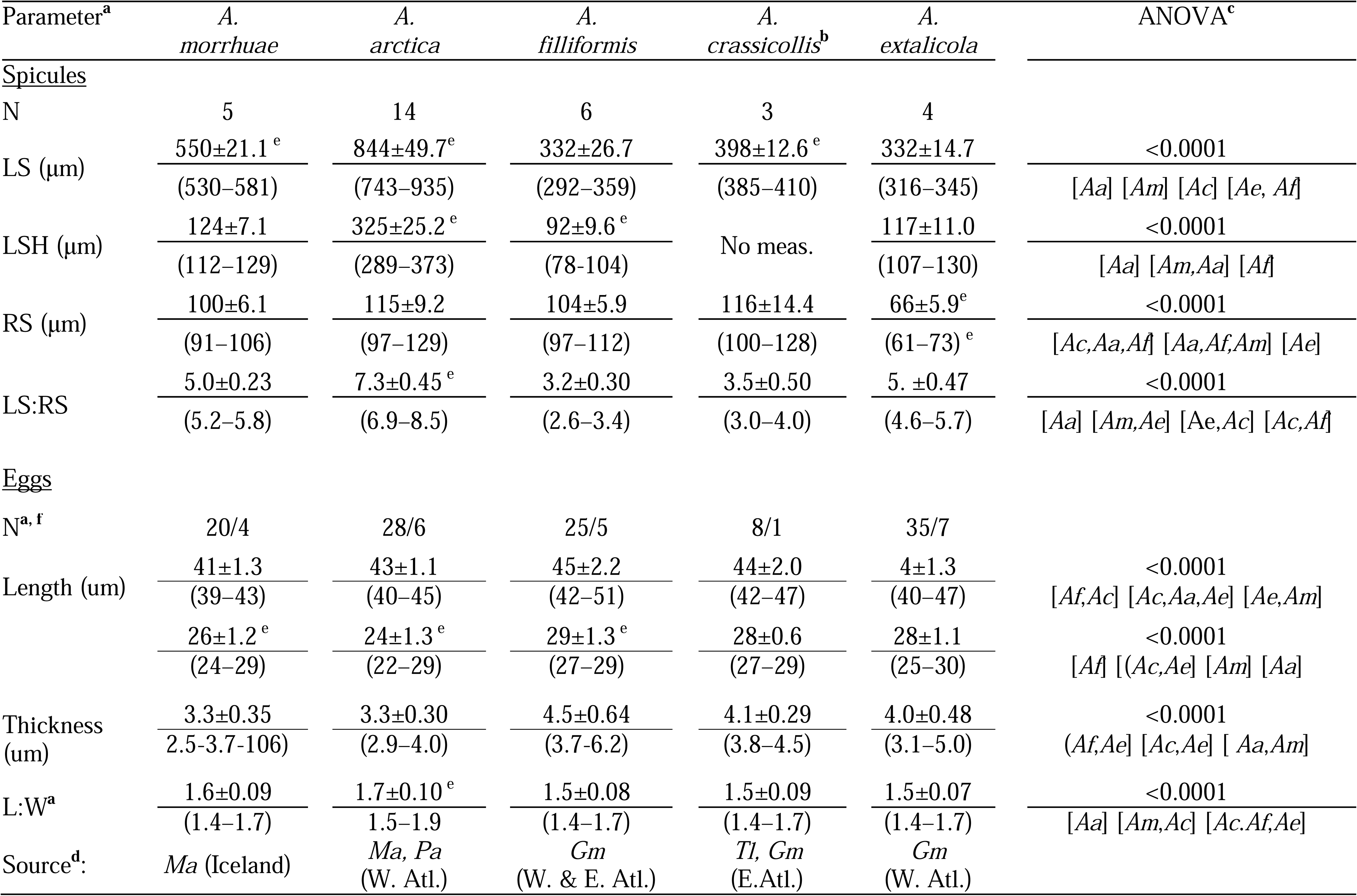
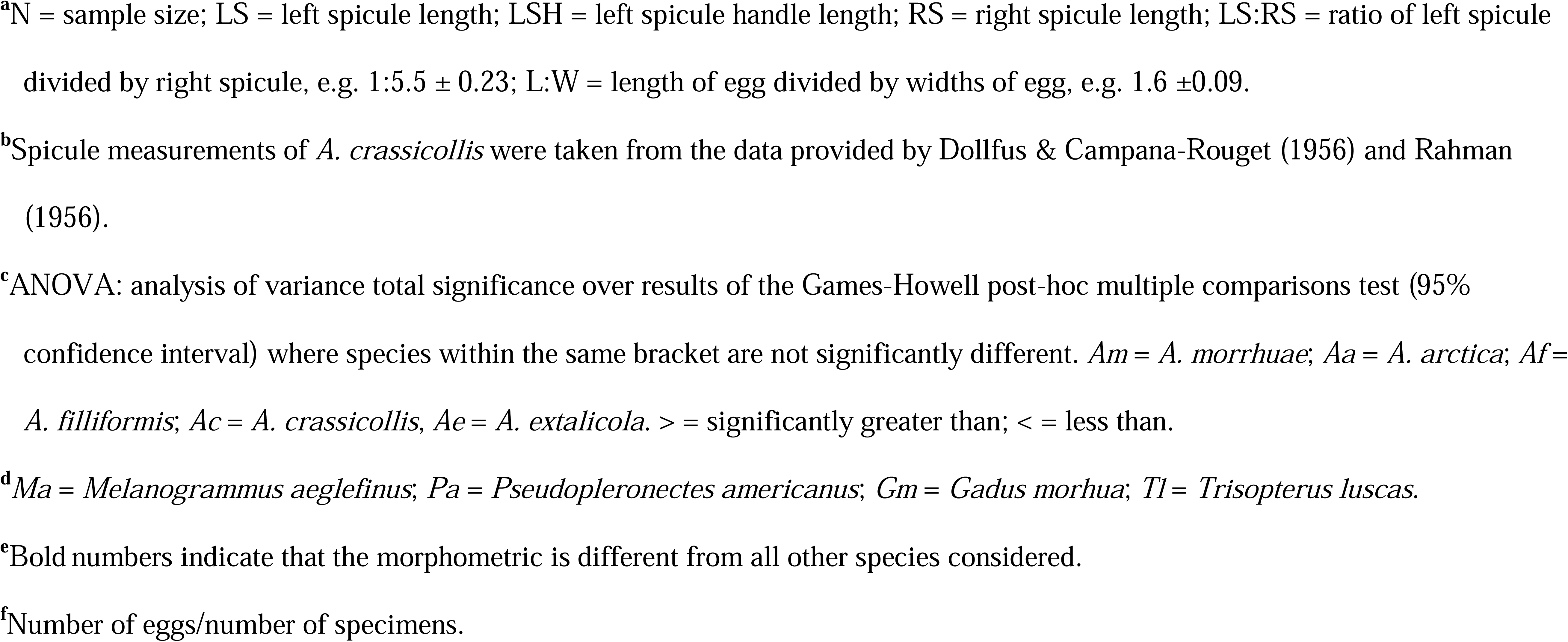
Detailed comparison of spicule and egg morphometrics of *Ascarophis* species from North Atlantic fishes based on measurements conducted in the present study.

| Parameter<sup>a</sup> | A.<br>*morruhae* | A.<br>*arctica* | A.<br>*filliformis* | A.<br>*crassicollis*<sup>b</sup> | A.<br>*extalicola* | ANOVA<sup>c</sup> |
| --- | --- | --- | --- | --- | --- | --- |
| <u>Spicules</u> |  |  |  |  |  |  |
| N | 5 | 14 | 6 | 3 | 4 |  |
| LS (µm) | 550±21.1<sup>e</sup><br>(530–581) | 844±49.7<sup>e</sup><br>(743–935) | 332±26.7<br>(292–359) | 398±12.6<sup>e</sup><br>(385–410) | 332±14.7<br>(316–345) | <0.0001<br>[Aa] [Am] [Ac] [Ae, Af] |
| LSH (µm) | 124±7.1<br>(112–129) | 325±25.2<sup>e</sup><br>(289–373) | 92±9.6<sup>e</sup><br>(78–104) | No meas. | 117±11.0<br>(107–130) | <0.0001<br>[Aa] [Am,Aa] [Af] |
| RS (µm) | 100±6.1<br>(91–106) | 115±9.2<br>(97–129) | 104±5.9<br>(97–112) | 116±14.4<br>(100–128) | 66±5.9<sup>e</sup><br>(61–73)<sup>e</sup> | <0.0001<br>[Ac,Aa,Af] [Aa,Af,Am] [Ae] |
| LS:RS | 5.0±0.23<br>(5.2–5.8) | 7.3±0.45<sup>e</sup><br>(6.9–8.5) | 3.2±0.30<br>(2.6–3.4) | 3.5±0.50<br>(3.0–4.0) | 5. ±0.47<br>(4.6–5.7) | <0.0001<br>[Aa] [Am,Ae] [Ae,Ac] [Ac,Af] |
| <u>Eggs</u> |  |  |  |  |  |  |
| N<sup>a, f</sup> | 20/4 | 28/6 | 25/5 | 8/1 | 35/7 |  |
| Length (um) | 41±1.3<br>(39–43) | 43±1.1<br>(40–45) | 45±2.2<br>(42–51) | 44±2.0<br>(42–47) | 4±1.3<br>(40–47) | <0.0001<br>[Af,Ac] [Ac,Aa,Ae] [Ae,Am] |
|  | 26±1.2<sup>e</sup><br>(24–29) | 24±1.3<sup>e</sup><br>(22–29) | 29±1.3<sup>e</sup><br>(27–29) | 28±0.6<br>(27–29) | 28±1.1<br>(25–30) | <0.0001<br>[Af] [(Ac,Ae) [Am] [Aa] |
| Thickness (um) | 3.3±0.35<br>(2.5–3.7–106) | 3.3±0.30<br>(2.9–4.0) | 4.5±0.64<br>(3.7–6.2) | 4.1±0.29<br>(3.8–4.5) | 4.0±0.48<br>(3.1–5.0) | <0.0001<br>(Af,Ae) [Ac,Ae] [ Aa,Am] |
| L:W<sup>a</sup> | 1.6±0.09<br>(1.4–1.7) | 1.7±0.10<sup>e</sup><br>(1.5–1.9) | 1.5±0.08<br>(1.4–1.7) | 1.5±0.09<br>(1.4–1.7) | 1.5±0.07<br>(1.4–1.7) | <0.0001<br>[Aa] [Am,Ac] [Ac,Af,Ae] |
| Source<sup>d</sup>: | Ma (Iceland) | Ma, Pa<br>(W. Atl.) | Gm<br>(W. & E. Atl.) | Tl, Gm<br>(E.Atl.) | Gm<br>(W. Atl.) |  |

<sup>a</sup>N = sample size; LS = left spicule length; LSH = left spicule handle length; RS = right spicule length; LS:RS = ratio of left spicule divided by right spicule, e.g. 1:5.5 ± 0.23; L:W = length of egg divided by widths of egg, e.g. 1.6 ± 0.09.
<sup>b</sup>Spicule measurements of *A. crassicollis* were taken from the data provided by Dollfus & Campana-Rouget (1956) and Rahman (1956).
<sup>c</sup>ANOVA: analysis of variance total significance over results of the Games-Howell post-hoc multiple comparisons test (95% confidence interval) where species within the same bracket are not significantly different. *Am* = *A. morrhuae*; *Aa* = *A. arctica*; *Af* = *A. filliformis*; *Ac* = *A. crassicollis*, *Ae* = *A. extalicola*. > = significantly greater than; < = less than.
<sup>d</sup>*Ma* = *Melanogrammus aeglefinus*; *Pa* = *Pseudopleuronectes americanus*; *Gm* = *Gadus morhua*; *Tl* = *Trisopterus luscus*.
<sup>e</sup>Bold numbers indicate that the morphometric is different from all other species considered.
<sup>f</sup>Number of eggs/number of specimens.

**Table 6.** Comparison of key morphological characteristics of *Ascarophis* species present in the North Atlantic and adjacent seas. Bold

|  | *A. morrhuae*<sup>a</sup> | *A. arctica*<sup>a</sup> | *A. filliformis*<sup>a</sup> | *A. crassicollis*<sup>a</sup> | *A. extalicola*<sup>a</sup> |
| --- | --- | --- | --- | --- | --- |
| Location | Stomach | Stomach | Stomach | Stomach | Rectum |
| Length (mm) | M<5<br>F<8 | M<8<br>F<17 | M<15<br>F<36 | M<4<br>F<8 | M<7<br>F<9 |
| Max. Width (µm) | M<80,<br>F<150 | M<105<br>F<160 | M<130<br>F<210 | M~95<br>F<170 | M<52<br>F<72 |
| Striations<sup>b</sup> | Very pronounced.<br>Well defined<br>beginning abruptly<br>at nerve ring.<br>Perpendicular to<br>long body axis | Pronounced. Well<br>defined beginning<br>gradually at junction of<br>muscular and glandular<br>esophagus. Angular to<br>long body axis. | Fine | Fine | Fine |
| Left spicule length<br>(µm)<sup>c</sup> | Long<br>(530–581) | Long<br>(743–935) | Short<br>(292–359) | Short<br>(385–410) | Short<br>(316–345) |
| Left spicule tip<sup>d</sup> | Straight, fine taper | Bent, inflated | Straight | Straight | Slightly<br>inflated |
| Eggs<sup>e</sup> | 2 broad filaments<br>at one pole,<br>sometimes a 3<sup>rd</sup><br>filament,<br>prominent knob | 2–4 broad & <14 fine<br>filaments at each pole | 2 broad<br>filaments at<br>one pole,<br>sometimes a<br>3<sup>rd</sup> filament,<br>prominent<br>knob | No filaments | 2–12<br>filaments<br>varying<br>diameter at<br>one pole |

**Table 7.**
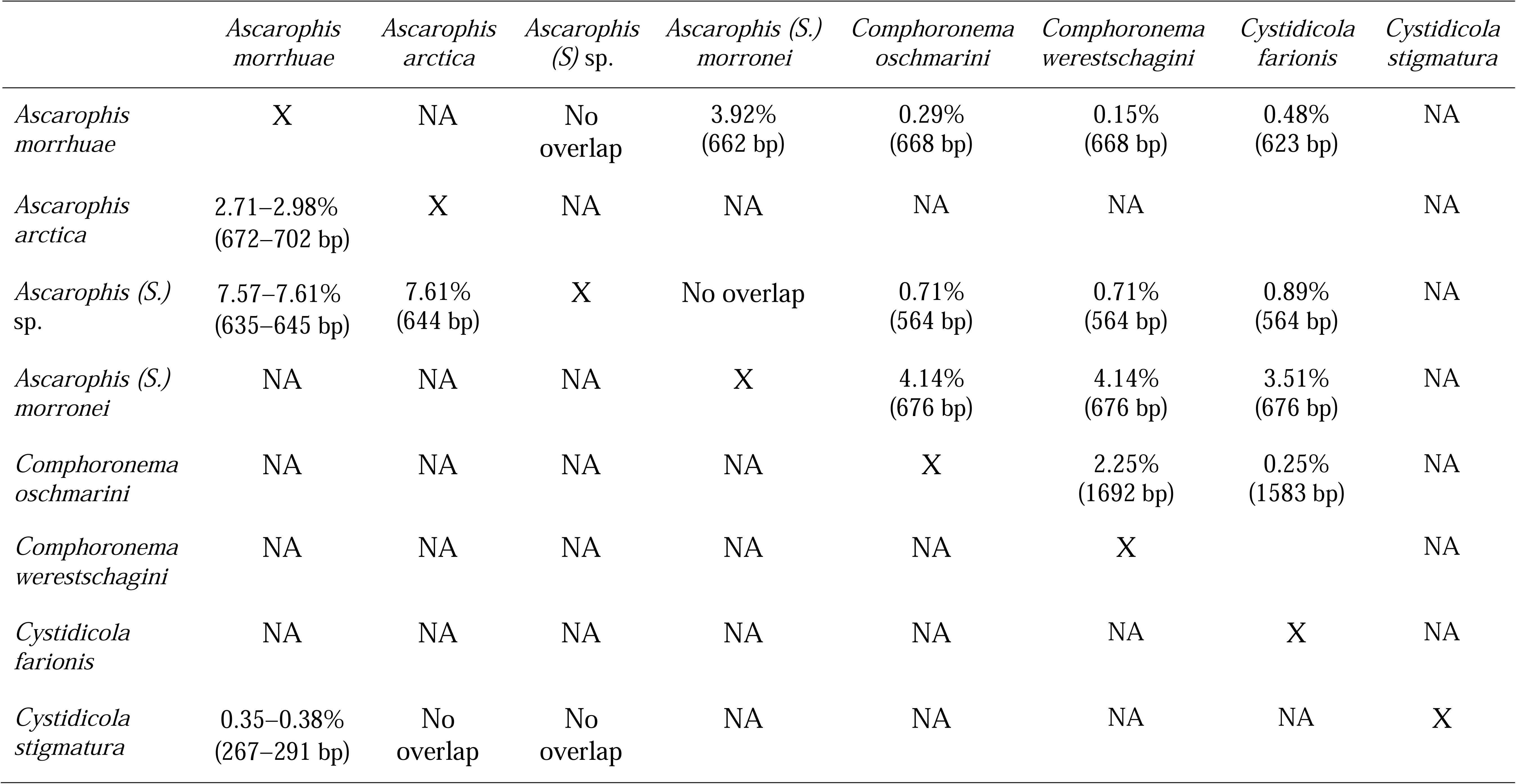
Pairwise genetic non-model corrected distances (%) between species present in the phylogenetic clade containing *Ascarophis morrhuae* Van Beneden, 1870. Distances in 28S rRNA gene portion below diagonal and 18S rRNA gene portion above diagonal with the length of overlapping region in parentheses.

|  | *Ascarophis*<br>*morrhuae* | *Ascarophis*<br>*arctica* | *Ascarophis*<br>*(S) sp.* | *Ascarophis (S.)*<br>*morronei* | *Comphoronema*<br>*oschmarini* | *Comphoronema*<br>*werestschagini* | *Cystidicola*<br>*farionis* | *Cystidicola*<br>*stigmatura* |
| --- | --- | --- | --- | --- | --- | --- | --- | --- |
| *Ascarophis*<br>*morrhuae* | X | NA | No<br>overlap | 3.92%<br>(662 bp) | 0.29%<br>(668 bp) | 0.15%<br>(668 bp) | 0.48%<br>(623 bp) | NA |
| *Ascarophis*<br>*arctica* | 2.71–2.98%<br>(672–702 bp) | X | NA | NA | NA | NA |  | NA |
| *Ascarophis (S.)*<br>*sp.* | 7.57–7.61%<br>(635–645 bp) | 7.61%<br>(644 bp) | X | No overlap | 0.71%<br>(564 bp) | 0.71%<br>(564 bp) | 0.89%<br>(564 bp) | NA |
| *Ascarophis (S.)*<br>*morronei* | NA | NA | NA | X | 4.14%<br>(676 bp) | 4.14%<br>(676 bp) | 3.51%<br>(676 bp) | NA |
| *Comphoronema*<br>*oschmarini* | NA | NA | NA | NA | X | 2.25%<br>(1692 bp) | 0.25%<br>(1583 bp) | NA |
| *Comphoronema*<br>*werestschagini* | NA | NA | NA | NA | NA | X |  | NA |
| *Cystidicola*<br>*farionis* | NA | NA | NA | NA | NA | NA | X | NA |
| *Cystidicola*<br>*stigmatura* | 0.35–0.38%<br>(267–291 bp) | No<br>overlap | No<br>overlap | NA | NA | NA | NA | X |

[Based on measurements of five male and six female worms and SEM of one male and two female worms].

GENERAL: Small, narrow nematodes; maximum width of body near posterior end. Cuticle thick, with overlapping transverse helical striations beginning abruptly near the nerve ring, diminishing posteriorly beginning at the level of junction of the glandular esophagus and intestine, with irregular lateral anastomoses. Cephalic end rounded, pseudolabia narrow with conical terminal protrusions. Oral opening elongate, demarcated by distinct submedian labia, and flap-like sublabia without indentations. Four submedian cephalic papillae and a pair of lateral amphids present. Buccal capsule long, cylindrical with funnel–shaped prostome in lateral view. Glandular esophagus four times longer than muscular. Nerve ring encircles muscular esophagus in anterior portion of muscular esophagus; excretory pore located posterior to level of nerve ring; small simple/unbifurcated deirids situated at about level of posterior end of buccal cavity.

MALE: Approximately 60% the length of female worms. Posterior end of body ventrally or spirally coiled, provided with narrow caudal alae, left ala extending slightly further anterior than right ala. Ten pairs of caudal papillae including four pedunculate pairs precloacal and six pairs postcloacal; five pedunculate and tenth pair minute, subventral and sessile. Caudal papillae pairs 1, 3, 6 digitiform, 2, 4, 5, and 7, more or less globose. Pair of small phasmids present just posterior to tenth pair of caudal papillae. Ventral *area rugosa* anterior to cloaca comprising eight to ten rows of tessellated cuticular ridges. Left spicule shaft approximately 23 percent of spicule length; tip of left spicule thin, narrow, without terminal inflation. Right spicule boat shaped. Left spicule approximately five times longer than right spicule. Tail conical, rounded. Testis originating in the area of the junction of the glandular esophagus and intestine, frequently reflexed, vas deferens junction with ejaculatory duct at approximately 70 percent of body from anterior end.

FEMALE: Worms without coiled tail. Amphidelphic; anterior ovary and uterus extending anteriorly to midpoint of the glandular esophagus; posterior ovary and frequently uterus extending posteriorly to level of anus. Uterus filled with numerous eggs, occupying a major part of the body. Vulva present at approximately 60 percent of body from anterior end, short vagina vera, vagina uterina directed posteriorly from vulva. Larvated eggs oval, thick–walled, two wide filaments supplemented frequently by a third and rarely by a fourth thin filament extending from a prominent knob on one pole; irregular papillae like structures on egg surface.

### Intraspecific morphological remarks

Besides *A. morrhuae* there have been eight species of *Ascarophis* reported from North Atlantic fishes, of which four are presently considered to be valid species. The four species not considered valid include: specimens of the North Pacific species *A. longispicula* Zhukov, 1960 reported from Baltic Atlantic cod (Studnicka, 1965) by Kirjušina & Vismanis (2007), are considered to be *A. arctica* by Appy (1981); *A. helix* Cobb, 1928, considered a *species inquirenda* (Dollfus & Campana-Rouget, 1956) was only briefly described from a female worm from the gills of a roughtail stingray, *Bathytoshia centroura* (Mitchill, 1815), from the Western Atlantic (Cobb, 1928); *A. baylisi* Dollfus & Campana-Rouget, 1956 from the streaked gurnard, *Chelidonichthys lastovize* (Bonnaterre, 1788) (as *Trigla lineata*), considered a synonym of *A. morrhuae* (Petter 1970; Rahman, 1966) (but see discussion below) and *A. skrjabini* (Layman, 1933) originally described from Eastern Asian freshwater/estuarine/anadromous fishes and subsequently reported from the fourhorn sculpin, *Myoxocephalus quadricornis* (Linnaeus, 1758), as *Cystidicola skrjabini* (Layman, 1933) in the Gulf of Finland (Dogiel and Rozova, 1941), from the eelpout, *Zoarces viviparous* (Linnaeus, 1758) and Atlantic cod in the Baltic (see Kirjušina and Vismanis, 2007), and considered a *species inquirenda* (Appy and Anderson, 1982; Skrjabin et al., 1967; Zhukov, 1960) and a composite taxon (Ko, 1986). Of the four recognized/valid species, *A. morrhuae* superficially is most similar to *A. arctica* in possessing a long, left spicule, having well defined striations, and filamented eggs. However, *A. morrhuae* differs from *A. arctica* in the following ways: it is shorter than *A. arctica* (Tables 3, 4); the pseudolabia of *A. morrhuae* are narrower compared to *A. arctica* as pictured by Fagerholm and Berland (1988) and Appy (1981); the tip of the spicule is tapered to a fine straight point in *A. morrhuae* while it is bent and inflated in *A. arctica* (Fig. 3F vs. Fig. 4C) (see also Appy, 1981; Fagerholm and Berland, 1988); the left spicule and spicule shaft (calomus) of *A. morrhuae* are shorter than those of *A. arctica* (Table 5); the eggs of *A. morrhuae* have two or sometimes three filaments emanating from a pronounced knob at one pole, while the eggs of *A. arctica* have many filaments at both poles (Figs. 1I, 2E, 3F, 4G vs. Figs. 4H). Based on present analysis, the eggs of *A. morrhuae* are on average wider than the eggs of *A. arctica* (Table 5); striations of *A. morrhuae* become pronounced just posterior to the nerve ring and most pronounced at the junction of the muscular and glandular esophagus, while those of *A. arctica* become pronounced at the level of the glandular esophagus and become most pronounced at the junction of the glandular esophagus and the intestine (Figs. 1B, 2B, 2C vs. Fig. 4A); and the striations are more perpendicular to the long body axis in *A. morrhuae* than are the striations of *A. arctica* (Fig. 2D vs. Fig. 4B). As mentioned above, *Ascarophis morrhuae* is consistently shorter than *A. arctica* (Tables 3, 4, 6).

**Figure 2.**
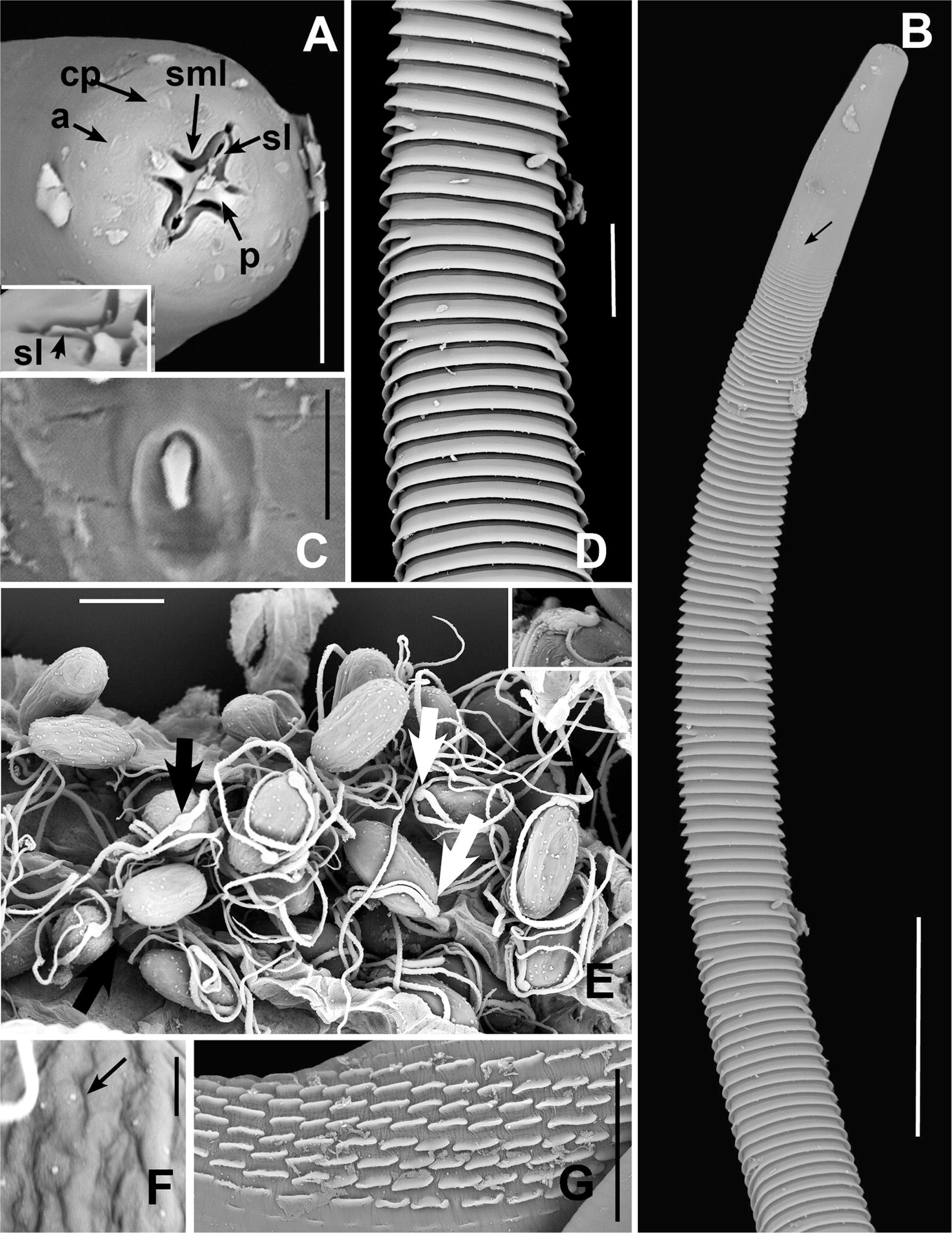
Scanning electron micrographs of *Ascarophis morrhuae* Van Beneden, 1870 from the the haddock, *Melanogrammus aeglefinus* (Linnaeus, 1758) from Iceland. A. Oral opening; inset-unnotched sublabia. Scale Bar: 10 μm. B. Anterior of worm showing striations relative to position of nerve ring (arrow). Scale Bar: 100 μm. C. Deirid. Scale Bar: 2.5 μm. D. Striations at approximate level of the junction of the muscular and glandular esophagus with anastomoses (arrow). Scale Bar: 25 μm. E. Eggs with 2 (black arrow) or 3 (white arrow) at one pole; inset: egg with 4 filaments. Scale Bar: 25 μm. F. Egg surface with papillae (arrow). Scale Bar: 2.5 μm. G. Area rugosa. Scale Bar: 20 μm. Abbreviations: a = amphid, cp = cephalic papilla, p = pseudolabia, sl = sublabia, sml = submedian labia.

**Figure 3.**
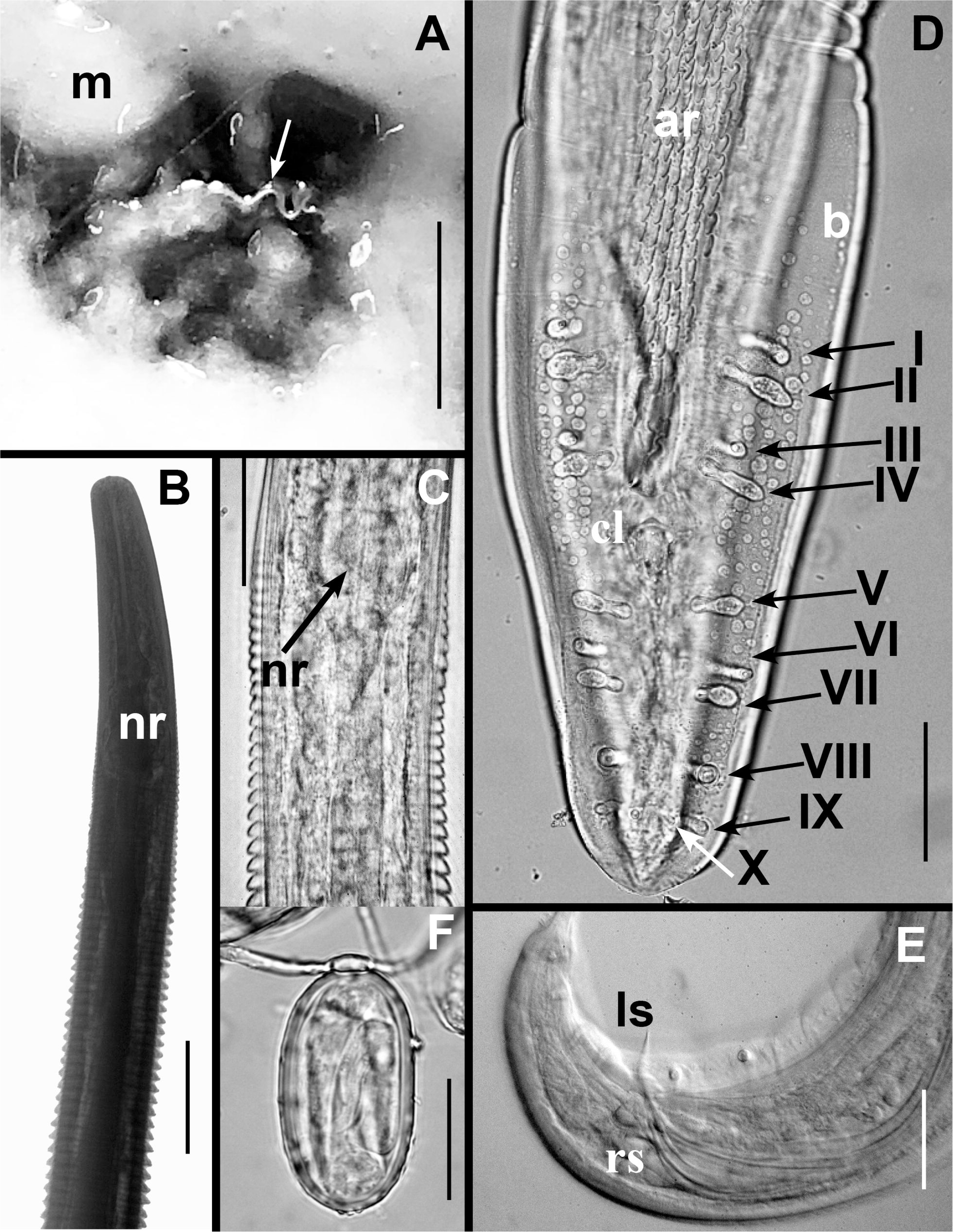
Light microscope photos of *Ascarophis morrhuae* Van Beneden, 1870 from the haddock, *Melanogrammus aeglefinus* (Linnaeus, 1758) from Iceland. A. worm present in mucus lining the pyloric stomach (white arrow). Scale Bar: 5 mm. B. Anterior end, striation profile relative to position of nerve ring. Scale Bar: 25 μm. C. Striation profile relative to position of nerve ring. Scale Bar: 25 μm. D. Ventral view of caudal end of male showing caudal papillae I – X. Scale Bar: 50 μm. E. Lateral view of caudal end of male. Scale Bar: 30 μm. F. Egg. Scale Bar: 25 μm. Abbreviations: ar = area rugosa, cl = cloaca, b = bursa, ls = left spicule, m = mucosa, nr = nerve ring, rs = right spicule.

**Figure 4.**
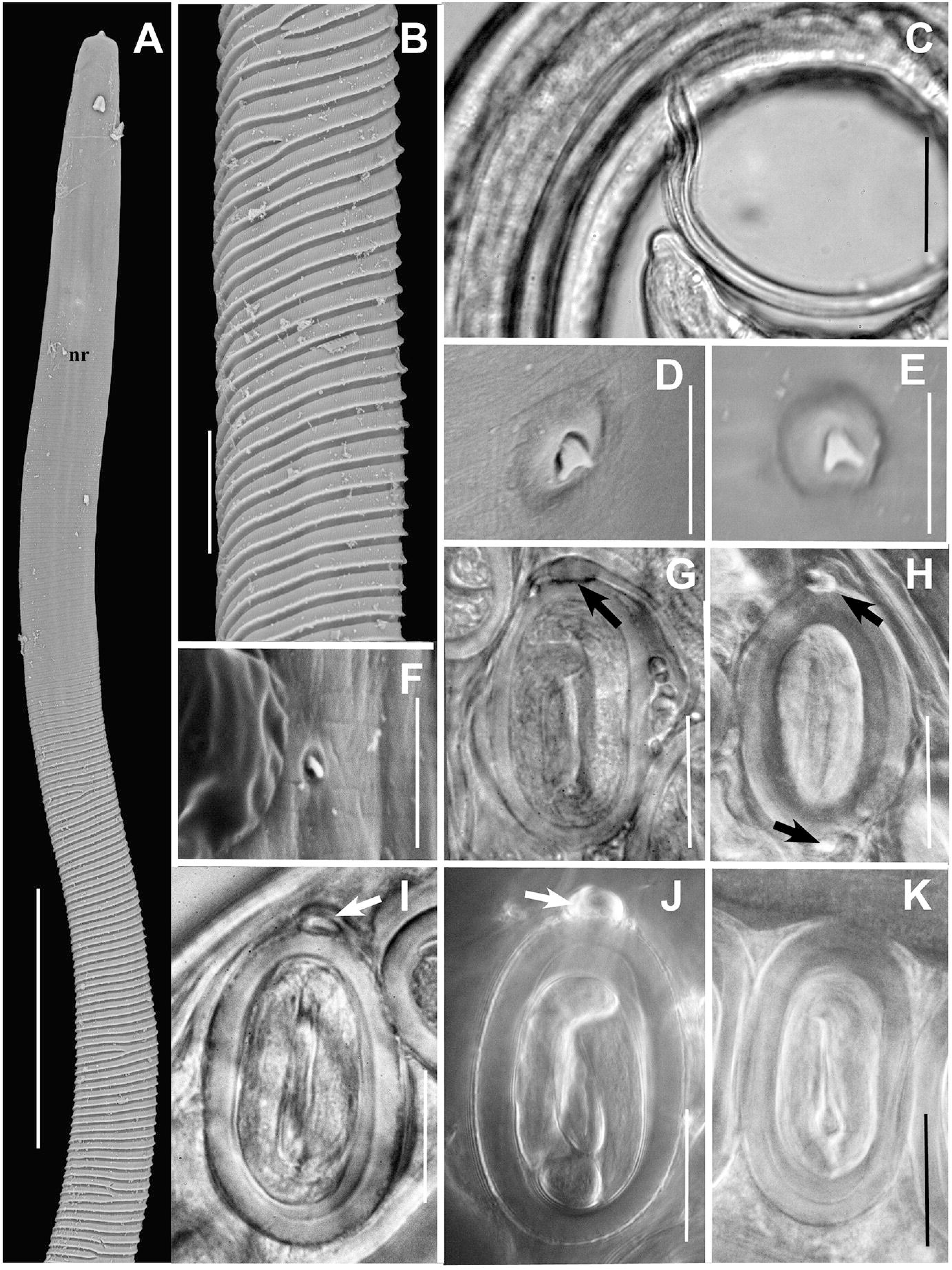
Electron micrographs and light microscope photos of *Ascarophis* species present in the North Atlantic. A. Anterior extremity of *A. arctica* Poljansky, 1952. Scale Bar: 100 μm. B. Striations of *A. arctica* at level of junction of glandular esophagus and intestine. Scale Bar: 25 μm. C. Tip of left spicule of *A. arctica*. Scale Bar: 30 μm. D – F: Deirids of *Ascarophis* species. Scale Bar: 5 μm: *A. arctica.* (D), *A. filliformis* Poljansky, 1952 (E), *A. extalicola* Appy, 1981 (F). G – K: Eggs of *Ascarophis* species. Scale Bar: 20 μm: of *A. morrhuae* Van Beneden, 1870 (G), *A. arctica (H)*, *A. extalicola (I)*, *A. filliformis (J)* and *A. crassicollis* Dollfus & Campana–Rouget 1956 (K) (polar knobs shown by arrows).

However, in specimens of *A. morrhuae* described by Ko (1986) from *Agonus cataphractus* (Linnaeus, 1758), both male and female worms are longer, and spicules are longer than specimens of *A. morrhuae* from the present study (Tables 3, 4) and longer than previous descriptions of *A. morrhuae* by Gordon (1951), Polyanskii (1952) and Srivastava (1966). In addition, specimens examined from the NHMUK on which Ko based his description (NHMUK 1962.462-481) have eggs with filaments at both poles of eggs (Fig. 4H), a long spicule bent at the tip and striations not becoming pronounced until the level of the muscular and glandular esophagus. *Ascarophis morrhuae* can be distinguished from the remaining three species from the North Atlantic by its pronounced striations, long left spicule, and through a combination of other features (Table 6). In addition, the deirids of *A. morrhuae* and *A. extalicola* are simple/unbifurcated while those of *A. arctica* and *A. filiformis* are bluntly bifed (Fig. 2C, 4F vs. Fig. 4D, 4E). The eggs of *A. morrhuae* are similar to those of *A. filiformis* in possessing two or three filaments emanating from a knob at one pole although the knob is much more prominent in *A. filiformis* and its eggs are wider on average than those of *A. morrhuae* (Table 5).

### Genetic characterization and phylogenetic placement

In total, eight specimens of *A. morrhuae* were genetically characterized with varying success across the selected genetic markers. The portion of 28S rDNA was generated for the three specimens of *A. morrhuae* originated from the Icelandic coast and a single one from the North Sea ranging from 994 bp to 1,025 bp with a single nucleotide difference (0.1%) between two of the specimens from the two localities. A single haplotype was retrieved for the portion of 18S rDNA generated for the two specimens of *A. morrhuae* originated from the Icelandic coast (660 bp to 668 bp) and five from the North Sea (663 bp to 681 bp), respectively. A single fragment of 674 bp of the COX1 mtDNA region was retrieved from the specimen of *A. morrhuae* collected from *G. morhua* from the North Sea.

Following the results of the molecular barcoding, the specimens of *A. morrhuae* from haddock collected in Icelandic waters and the North Sea are identical while they are distinct from all other cystidicolids present in the phylogenetic clade sequenced to date (Table 7). This also includes the other two sequenced species of *Ascarophis* (i.e., *A. (Ascarophis) arctica* and *A. (Similascarophis) morronei* following Muñoz et al. (2004)). *Ascarophis morrhuae* is present in a well-supported clade consisting of representatives of three other genera *Comephornema*, *Capillospirura* and *Cystidicola* with *A.* (*Dentiascarophis*) *adioryx* Machida, 1981 being part of a different clade. Phylogenetic relationships among the species of *Ascarophis* in the cystidicolid clade, *sensu lato*, remain unresolved (Fig. 5).

**Figure 5.**
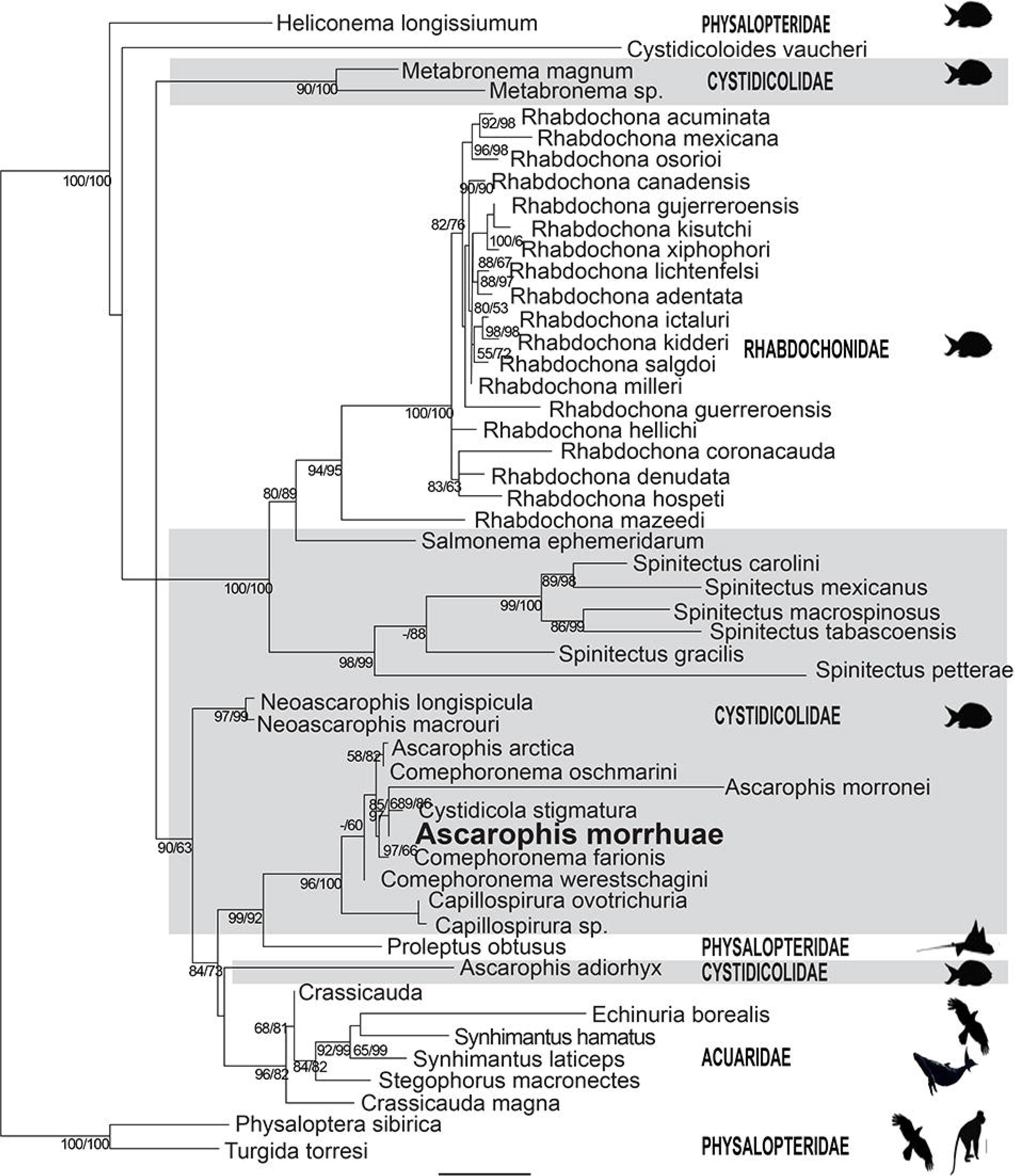
Phylogenetic relationships between *Ascarophis morrhuae* Van Beneden, 1870 and other cystidicolid species based on concatenated portions of the 18S and 28S rRNA gene. Support values are presented as Ultrafast bootstrap values/SH-aLRT/Bayesian posterior probabilities. The scale bar represents the estimated number of substitutions per site. Shaded area represents species presently placed in the Cystidicolidae. Icons represent host groups infected including bony fishes, elasmobranchs, birds and mammals. Scale Bar: 0.04

## Discussion

After 155 years since the original brief description of Van Beneden (1870), the morphology of the enigmatic species, *A. morrhuae*, is fully redescribed using standard morphological techniques, scanning electron microscopy and genetic sequencing of newly collected material from Icelandic haddock and the type host, the Atlantic cod from the North Sea. This study differentiates *A. morrhuae* from North Atlantic species of *Ascarophis*, identifies the Eastern Atlantic, as far west as Iceland, and adjoining seas as the likely geographic distribution of *A. morrhuae*, and the haddock as a preferred host. Importantly, these findings allow establishment of neotypes and neogenotypes of one of the earliest described and basal cystidicolids, which provide a genetic and taxonomic reference for subsequent study of spiromorph nematodes.

The taxonomic history of *A. morrhuae,* following its brief description by Van Beneden (1870), has been provided by Dollfus (1953) and Skrjabin et al. (1967). Specimens of *A. morrhuae* collected during the present study conform morphologically with previous characterizations that can be ascribed with certainty to *A. morrhuae* Van Beneden, 1870 (Nicoll, 1907 as *Ascaropsis morrhuae* Power & Sedgwick, 1880 per Dollfus, 1953; Gordon, 1951; Polyanskii, 1952, Srivastava 1966; Van Beneden, 1870). The original two–sentence description^2^ [^2^*Ascarophis morrhuæ*, sp. nov. **^9^**, pl. III, fig. 11 (van Beneden, 1870): Ce ver est extrêmement remarquable, d’abord par sa petite taille et ensuite par la forme effilée de l’extrémité céphalique. Toute la surface du corps est finement striée. Les oeufs se distinguent de tous les autres par les filaments qui garnissent un des pôles (van Beneden, 1870).] and drawings of Van Beneden (1870) from a single female worm from an Atlantic cod, identified eggs with two prominent filaments and very anterior portion of body with distinct annulations (Van Beneden, 1870: p. 56 & Plate III, Fig. 11a, b). Subsequent to that, Nicoll (1907), Gordon (1951) and Polyanskii (1952) provided descriptions, including drawings, which are consistent with Van Beneden’s (1870) description having prominent striations anteriorly and eggs with two filaments emanating from a prominent knob on one pole (Fig. 11 in Van Beneden, 1870). In addition, Gordon (1951), Polyanskii (1952) and Srivastava (1966) described/illustrated the tip of the left spicule as tapering to a fine point consistent with present material. Polyanskii (1952) provided the first relatively complete morphometric description of *A. morrhuae* based on a number of specimens from both cod and haddock collected in the Barents Sea, and which conforms to specimens described herein.

However other specimens have been attributed incorrectly to *A. morrhuae*. In 1933, Baylis identified two large female worms (11.2 mm and 13.2 mm) from *C. lastovize.* These specimens were later referred to a new species, *A. baylisi* Dollfus and Campana-Rouget, 1956 due their large size, lack of filamented eggs and host (Dollfus and Campana-Rouget, 1956). Subsequently, Rahman (1966) synonymized *A. baylisi* with *A. morrhuae* and Petter (1970) concurred, believing that the length was a variable character, the lack of filaments was due to the immaturity of the eggs (i.e., non larvated) and both worms were striated. Examination of the Baylis specimens present in the NHMUK (1932.12.6 2/3) shows they differ from *A. morrhuae* in their large size, and lack of pronounced striations in the area of the nerve ring. It is also notable that the eggs of the Baylis specimens are immature at 13 mm while specimens of *A. morhuae,* 5 mm in length, are packed with larvated eggs. While the size and striations of the Baylis worms conform with *A. arctica*, additional material including mature females and males from *C. lastoviza* are needed to confirm this identification and determine the egg morphology. As a result, the Baylis material is reinstated as *A. baylisi* as a *species inquirienda*.

The use of egg filaments in the identification of species of *Ascarophis* is valuable (Ferrer et al., 2005) with some precautions. The presence of filaments at poles should be confirmed by examining a large number of eggs and viewing larvated eggs in lateral view in utero, since filaments may detach from the egg during fixation and manipulation of worms for study; this probability is compounded in species with filaments at both poles. While Moravec and Nagasawa (2018) recently provided a scanning electron micrograph of an egg of *A arctica* with filament at only one pole (vs filaments at both poles), this should be considered a rare occurrence. Likewise, the number of filaments at a pole is a reliable character within reason.

*Ascarophis morrhuae* and *A. filliformis* both were described as having two filaments at one pole while more recent studies using SEM have shown that eggs with a third and rarely four filaments in the former (present study) and three in the latter (Ko, 1986). However, in both cases, the third and fourth filaments are thin. Special caution should be taken with describing species based on filament number where there are many filaments at one or both poles, since their enumeration is difficult, and filament number may increase as the eggs are formed in the uterus (Ferrer et al., 2005). In any regard, examination of many larvated eggs both in utero and extra utero should be carried out as part of species descriptions/identifications. As a further precaution, two cystidicolids, i.e., *Cystidicola stigmatura* (Leidy, 1886) and *C. farionis* Fischer, 1898, which have distinct egg morphologies, have identical rDNA sequences (ITS-1, 5.8S and 28S(D3), while two morphologically identical isolates within both species have four different nucleotide positions in the ITS-2 sequences (Miscampbell et al. 2004).

Some historic reports of *A. morrhuae* do not conform with findings in the present study and those of Nicoll (1907), Gordon (1951) and Polyanskii (1952). For example, Punt (1947), Ko (1986), Petter (1970) and Køie et al. (2008) depict/describe worms that are larger than *A. morrhuae*, have filaments at both egg poles, or lack pronounced striations at the level of the nerve ring. In his preliminary review of *Ascarophis*, Ko (1986) based his measurements of *A. morrhuae* on specimens from hooknose, *Agonus catafractus* (Linnaeus, 1758) present in the NHMUK but noted that there was size variation in his own description and in previous descriptions of *A. morrhuae.* Examination of material from *A. catafractus* present in the NHMUK, including material that formed the basis of Ko’s morphometric description included specimens which possessed eggs with filaments at both poles, a left bent spicule in the single male present, and striations becoming prominent at the junction of the muscular and glandular esophagus all features consistent with *A. arctica* and not *A. morrhuae*. *Ascarophis arctica* was also reported from *A. catafractus* in the North Sea by Klimpel et al. (2003). However, worms from *M. aeglefinus* utilized by Ko (1986) for SEM are likely *A. morrhuae* since all specimens from *M. aeglefinus* present in the NHMUK, where Ko obtained his specimens, are *A. morrhuae*.

While specimens described herein can confidently be identified as *A. morrhuae,* the morphological description is based on worms not taken from the type host, Atlantic cod, or presumed type locality (off Belgium) as recommended by Dollfus (1953) and Moravec and Nagasawa (2018). Attempts to obtain Atlantic cod that could be attributed to Belgium from Ostend, Belgian fish markets or from trawls carried out as part of this study along the coast of Belgium were not successful. While no *A. morrhuae* were present in Atlantic cod in Belgian waters, the sequences of both portions of 18S and 28S rDNA region of the worms from Iceland show 100% and 99.9% similarity, respectively, with specimens collected from Atlantic cod and haddock originated from the North Sea off Scotland. It also appears that while cod is the type host, haddock is a more common host of this species (Gordon, 1951; Nicoll, 1907; Polyanskii, 1952). Polyanskii (1966) found 43% of 76 haddock and only 4.4% of 90 cod infected with *A. morrhuae* (Polyanskii’s Table 16, p. 118 and Table 15 p. 114 respectively). All five haddock collected in the present study from Iceland were infected while two cod were uninfected. All specimens of *Ascarophis* from haddock in the NHMUK are *A. morrhuae.* While strict host specificity is not common among cystidicolids (see Moravec et al., 2024), it appears that *A. morrhuae* predominantly infects haddock. As a practical matter it may never be possible to find *A. morrhuae* from the type host/locality due to it being less common in Atlantic cod, and the significant decline in cod in the southern North Sea due to overexploitation of stocks and/or warming water due to climate change (Engelhard et al., 2014; Froese & Quaas, 2012), although cod have been reported associated with a wind farm in Belgian waters (Reubens et al., 2010). Based on these factors, specimens of *A. morrhuae* from Icelandic cod are herein established as neotypes and neogenotypes.

The geographic distribution of *A. morrhuae* based on confirmed reports, is restricted to the Eastern North Atlantic and its adjacent seas (e.g. North Sea and Barents Sea) as far West as Iceland. *Ascarophis morrhuae* reported in other geographic regions off Portugal (Rodrigues et al., 1973), Greenland (Koi et al., 2008) and Western North Atlantic (Hanek & Threlfall, 1970) and others (see listing in species description), could not be confirmed as belonging to *A. morrhuae* because specimens were not deposited in a museum, and/or were not made available for study. At least some worms attributed to *A. morrhuae* in the Western Atlantic/Greenland appear to be more similar to *A. arctica.* For example, Koi et al. (2008) describes *A. morrhuae* from Arctic cod, *Arctogadus glacialis* (Peters, 1872) from off Greenland as striated worms with filaments at both egg poles, characters more attributed to *A. arctica*. It appears that *A. morrhuae* is absent in the Northwestern Atlantic; the *Ascarophis* species found in intensive studies of haddock (Scott, 1981), Atlantic cod and other fishes (Appy, 1981; Arthur & Albert, 1993) in the Western Atlantic (i.e., Bay of Fundy, Scotian Shelf and Gulf of St. Lawrence) are *A. arctica*, *A. filiformis* or *A. extalicola*. Two of the five recognized species in the North Atlantic, *A. arctica* and *A. filiformis* (Blaylock et al, 1997; Ko, 1986; Zhukov, 1960) have also been reported from the Northern Pacific (Blaylock et al., 1997; Ko, 1986; Moravec & Nagasawa, 2018; Zhukov, 1960).

The morphology of *A. morrhuae* described herein is typical of the generally accepted diagnosis of *Ascarophis* (Ko, 1986) and the subgenus *A*. (*Ascarophis*) of Moravec & Nagasawa (2018). In particular, features of *A. morrhuae* visualized for the first time using SEM and DIC include a dorsoventrally elongate/oval oral opening, narrow pseudolabia with conical terminal protrusion anteriorly and connected posteriorly to anterior margin of the buccal capsule, four submedian labia, four flap-like sublabia four pairs of precloacal caudal papillae and six pairs of postcloacal caudal papillae in male worms. The other two subgenera *Similascarophis* and *Dentiascarophis* do not coincide with the generally accepted generic description of Ko (1986) since they lack submedian labia and *Dentiascarophis* has two protrusions/denticles associated with the base of the pseudolabia. However, modification of the generic diagnosis of *Ascarophis* is premature pending additional genetic studies on additional species among the subgenera of *Ascarophis* particularly since Sokolov et al. suggest *Dentiascarophis* should be elevated to generic status on genetic grounds.

The form of the deirids as viewed with SEM may be a useful taxonomic feature in cystidicolids (Moravec et al., 2007) being either simple, bifed or trifed (Beveridge and Moravec, 2020). In the present study deirid morphology of three species of *Ascarophis* (i.e., *A. morrhuae*, *A. filiformis* and *A. extalicola*) are added to the 11 species examined with SEM (Moravec, 2024). Of these 14 species, four have simple deirids while the remaining 10 species have bifurcated deirids, although there is considerable variation within these general morphological groups. The simple deirids of *A. morrhuae* are slightly curved and rounded, while those of *A*. *mexicana* are stick-like (Moravec & González-Solis, 2007). Among species of *Ascarophis*, with bifed deirids, the furcae may be narrow extending from an equally narrow base (e.g., *A. (Similascarophis) richeri* Moravec & Jiustine, 2007 and *A. arctica*) (Moravec and Justine, 2007 and Moravec and Nagasawa, 2018, respectively,) while in *A. (Dentiascarophis) adioryx*, the furcae appear as long conical structures surficially unattached at the base (Moravec et al., 2009) or *A.* (*Ascarophis*) *nasonis* Moravec & Justine, 2020, where the furcae are separated at their base by tissue (Moravec & Juistine, 2020). Also, there is no apparent correlation between deirid morphology and patterns of oral morphology. For example, within the *Ascarophis* subgenera (Moravec and Justine, 2009), *A.* (*Ascarophis*) contains species with simple deirids (e.g, *A. morrhuae*) and species with bifurcated deirids (e.g. *A. arctica*) and in the case of the examples cited, the oral morphology is nearly identical. The two remaining subgenera, *A*. (*Similascarophis*) and *A*. (*Dentiascarophis*) have species with only bifed deirids. This variation in deirid morphology is apparent within and between other genera of cystidicolids (e.g. *Neoascarophis* (Rossin et al., 2012 cf. Pereira et al., 2012) and *Comephoronema* (Moravec et al., 2007 cf. Pereira et al., 2014). While deirids are clearly a useful morphological tool in distinguishing cystidicolid species, based on present understanding, there is no apparent correlation between the morphology of deirids and oral structure within and between cystidicolid genera.

While the structure of the small oral structure of cystidicolid nematodes are traditionally generic features, their importance has been questioned by a number of authors (Ferrar et al., 2005; Moravec, 2007; Moravec & González-Solís, 2007; Moravec & Justine, 2009). This observation is due in part to polyphyly appearing within the Cystidicolidae and between the Cystidicolidae and the related Acuaridae and Physalopteridae as visible in the phylogenetic reconstruction following the results of the previous studies of Aguilar-Aguilar et al., (2019), Choudhury and Nadler, (2018), Černotíková et al., (2011), Jabbar et al., (2015), Sokolov et al. (2019) and Vidal et al., (2016). Any phylogenetic reconstruction of Cystidicolidae is hampered by the limited number of species for which sequence data are available. As shown in Table 2, only a small subset of species has sequence information for both genetic markers, 18S rDNA and 28S rDNA. This limitation is further compounded by the fact that the sequenced regions of these markers often do not overlap among species (see Table 7), which substantially reduces the amount of comparable data and consequently prevents strong support for the inferred topologies.

However, despite these limitations, *A. morrhuae* is part of a well-supported clade which includes *A. (Ascarophis) arctica* and *A. (Similascarophis) morronei*, and also includes the cystidicolid genera *Cystidicola*, *Comephoronema* and *Capillospirura*. while excluding *A. (Dentiascarophis) adioryx*. Thus the present study supports the suggestion of Sokolov et al. (2019) that the subgenus *Dentiascarophis* be elevated to genus.

Cystidicolids are predominantly parasites of the stomach, but their location within the host gut/stomach is variable. Appy (1981) found that *A. filliformis* was predominantly found in cardiac stomach, frequently among food items of the Atlantic cod, *A. arctica* was predominantly present in the pyloric stomach or occasionally among food items and *A. extalicola* with its capillarid-like body is only present among the villi of the rectum. Ko (1985) found *A. minuta* Ko, 1985 within the external muscularis of the stomach wall. Based on the present study, *A. morrhuae* is found in the mucus of the pyloric stomach. *Ascarophis morrhuae* is unique among cystidicolids in having highly developed transverse (vs angular) striations beginning abruptly at the area of the nerve ring, which may be an adaptation for moving through the dense mucus of the pyloric stomach. Cuticular structure of overlapping rings in criconematid nematodes allows them to greatly replace typical serpentine movement by extending and contracting the length of their bodies (Maggenti, 1981) and this may be the same for *A. morrhuae*. Oral and cuticular adaptations to microhabitat may, through convergent evolution, be responsible in part for the discord occurring between nematode families seen in phylogenetic studies. This may become clearer with publication of genetic information on the many genera and species only known from morphological study.

## Acknowledgements

Funding for travel and lodging of the senior author was made possible through an Incoming Mobility Grant from the Special Research Fund of Hasselt University (BOF24KV14). The data generation was funded by AfroWetMaP project of the Belgian Federal Science Policy Office (4255-FED-tWIN-G3 program, Prf-2022-049); infrastructure was funded by EMBRC Belgium – FWO project GOH3817N and DiSSCo Flanders 2.0 (R-15690). Help in collecting samples in Belgium was provided by Maarten Van Steenberge and Nathan Vranken (Hasselt University, Royal Belgian Institute of Natural Sciences). Cod and stomach samples from the North Sea were collected by Jim Drewery and other Marien Scotland staff aboard FRS Scotia. Technical help at the Hasselt University, Campus Diepenbeek was provided by Natascha Steffanie, Ria Vanderspikken, Tiziana Gobbin, Martia Topić and Miriam Shigoley. Genetic sequences of *A. (S.) morronei* were provided by Dr. Shana Goffredi, Occidental College, Department of Biology, Los Angeles, CA, USA.

## Authorship contribution statement

**Ralph G. Appy:** conceived and designed the project with support from MPMV and NK, collected specimens from haddock, analyzed the data, prepared the Figures and Tables and wrote the manuscript.

**Maarten P. M. Vanhove:** provided lab facilities at Hasselt University, helped with funding acquisition and provided technical review of the manuscript.

**Ken MacKenzie:** provided specimens collected from haddock in the North Sea and reviewed the manuscript.

**Jesús S. Hernández–Orts:** provided lab facilities and specimens from the cystidicolid collection at the NHMUK and reviewed the manuscript.

**Nikol Kmentová:** Helped conceive and design the project, supervised all aspects of the project, conducted the genetic analysis and reviewed and helped write the manuscript.

## Ethical approval

Parasite specimens were obtained from dead fishes obtained as bycatch from research vessels and/or Belgian commercial and retail fish purveyors.

## Disclosure statement

No potential conflict of interest was reported by the authors.

## Specimen availability

The neoholotype and neoallotype of *A. morrhuae* and two paratypes are deposited in the NHMUK. Two neoparatypes have been deposited in each of the USNM and HU. Voucher specimens of *A. arctica*, *A. filliformis*, and *A. extalicola* have been deposited in the USNM. Accession numbers are provided in Table 1.

## Data availability statement

The newly generated sequences are deposited in the GenBank database under the accession numbers provided in Table 2.

## Notes

### Competing Interest Statement

The authors have declared no competing interest.

## References

Aguilar–Aguilar, R. G., Ruiz–Campos, S., Martorelli M. M., Montesand A., & Martinez–Aquino, A. (2019) A new species of *Ascarophis* (Nematoda: Cystidicolidae) parasitizing *Clinocottus analis* (Pisces: Cottidae) from Baja California, Mexico Journal of Parasitology 105, 524–532.

Anderson, R. C. (2000) Nematode Parasites of Vertebrates: Their Development and Transmission. 2^nd^ Edition. CABI Publishing, Wallingford.

Anderson R. C., Chabaud A. G., & Willmott S (Eds.) (2009) Keys to the Nematode Parasites of Vertebrates. Archival Volume. CABI, Oxfordshire, UK 463 pp.

Appy, R. G., (1981) Species of *Ascarophis* Van Beneden, 1879 (Nematoda: Cystidicolidae) in North Atlantic fishes. Canadian Journal of Zoology 59, 2193–2205.

Appy, R. G., & Anderson, R. C. (1982) The genus *Capillospirura* Skrjabin, 1924 (Nematoda: Cystidicolidae) of sturgeons. Canadian Journal of Zoology 60, 194–202. 10.1139/z82-027.

Appy, R. G., & Butterworth, E. W. (2011) Development of *Ascarophis* sp. (Nematoda: Cystidicolidae) to maturity in *Gammarus deubeni* (Amphipoda). Journal of Parasitology 97, 1035–1048. 10.1645/GE-2878.1.

Arthur J. R., & Albert, E. (1993) A survey of the parasites of Greenland halibut (*Reinhardtius hippglossoides*) caught off Atlantic Canada, with notes on their zoogeography in this fish. Canadian Journal of Zoology 72, 765–778. 10.1139/z94-103.

Baylis, H. A. (1933) The nematode genus *Ascarophis* Van Beneden. Annals and Magazine of Natural History 11, 112–117.

Baylis, H. A. (1939) Further records of parasitic worms from British vertebrates. The Annals and Magazine of Natural History 23, 473–498.

Berland, B. (1961) Nematodes from some Norwegian marine fishes. Sarsia 2, 1–50.

Beveridge, I., & Moravec, F. (2020) *Ascarophisnema hoiae* n. sp. (Nematoda: Cystidicolidae), from the stomach of the trumpeter whiting, *Sillago maculata* Quoy & Gaimard (Perciformes: Sillaginidae) from Moreton Bay, Queensland, Australia. Systematic Parasitology 97, 297–304. 10.1007/s11230-020-09910-y.

Blaylock, R. B., Holmes, J. C., & Margolis L. (1997) The parasites of Pacific halibut (*Hippoglossus stenolepis*) in the eastern North Pacific: host-level influences. Canadian Journal of Zoology 76, 536–547. 10.1139/z97-214.

Černotíková, E., Horák, A., & Moravec, F. (2011) Phylogenetic relationships of some spirurine nematodes (Nematoda: Chromadorea: Rhabditida: Spirurina) parasitic in fishes inferred from SSU rRNA gene sequences. Folia Parasitologica 58, 135–148. 10.14411/fp.2011.013.

Cheethan, C. & Fives, J. M, (1990) The biology and parasites of the butterfish *Pholis gunnellus* (Linnaeus, 1758) in the Galway Bay area. Proceedings of the Royal Irish Academy, Section B: Biological, Geological and Chemical Science 90B, 127–149.

Choudhury, A., & Nadler, S. A. (2018) Phylogenetic relationships of spiruromorph nematodes (Spirurina: Spiruromorpha) in North American freshwater fishes. Journal of Parasitology 104, 496–504. 10.1645/17-195.

Cobb, N. A. (1928) The screw-nemas, *Ascarophis* Van Beneden, 1871; parasites of codfish, haddock and other fishes. Journal of the Washington Academy of Sciences 18, 96–102.

Dogiel, V. A., & Rozova A. (1941) Parasitic fauna of *Myoxocephalus quadricornis* in various areas of its distribution. Uchenye Zapiski Leningrudskogo Ordena Lenina Ciosudarstvennogo Universiteta 74, 14–19.

Dollfus, R. P. (1953) Parasites Animauix de la Morue Atlanto–Arctique. Encyclopédie Biologique Paris XLIII, 423 pp.

Dollfus, R. P., & Campana–Rouget, Y. (1956) Une nouvelle espeće d’*Ascarophis* (Nematoda, Spirurinae) chez *Gadus luscus* L. Révision de genre. Annales de Parasitologie Humaine et Comparée 31, 385–404.

Edgar, R. C. (2004). MUSCLE: Multiple sequence alignment with high accuracy and high throughput. Nucleic acids research 32, 1792–1797. doi: 10.1093/nar/gkh340.

Engelhard GH, Righton DA and Pinnegar JK (2014) Climate change and fishing: a century of shifting distribution in North Sea cod. Global Change Biology 20, 2473–2483. 10.1111/gcb.12513.

Fagerholm, H–P., & Berland, B. (1988) Description of *Ascarophis arctica* Poljanskyk 1952 (Nematoda: Cystidicolidae) in Baltic Sea fishes. Systematic Parasitology 11, 151–158. 10.1007/BF00012265.

Fagerholm H–P and Butterworth EW (1988) *Ascarophis* sp. (Nematoda: Spirurida) attaining sexual maturity in *Gammarus* spp. Crustacea 12, 123–139. 10.1007/BF00000147.

Ferrer E., Aznar, F. J., Balbuena, J. A., Kostadinova, A., Raga, J. A., & Moravec, F. (2005) A new cystidicolid nematode from *Mullus surmuletus* (Perciformes: Mullidae) from the Western Mediterranean. Journal of Parasitology 91, 335–344. 10.1645/GE-366R.

Froese, R., & Quoos, M. (2012) Mismanagement of the North Sea cod by the European Council. Ocean & Coastal Management 70, 54–58. 10.1016/j.ocecoaman.2012.04.005.

Gordon, A. R. (1951) On the male of *Ascarophis morrhuae* Van Beneden. Parasitology 41, 261–263.

Gaevskaya, A. V., & Umnova, B. A. (1977) On the parasite fauna of the principal commercial fishes of the northwest Atlantic. Biologiya Morya (Vladivostok) 4, 40–48.

Guindon, S., Dugayard, J-F., Lwefort, V., Anisimova, M., Hordijk, W., & Gascuel, O. (2010) New algorithms and methods to estimate maximum-likelihood phylogenies: assessing the performance of PhyML 3.0. Systematic Biology 59, 307–321. 10.1093/sysbio/syq010.

Hoang, D. T., Cheemor, O., Van Haeselar, A., Minh, B. K., & Vinh, L. S. (2018) Improving the ultrafast bootstrap extrapolation. Molecular Biology and Evolution 35, 518–522. 10.1093/molbev/msx281.

Hanek, G., & Threlfall, W. (1970) Parasites of the three–spine stickleback (*Gasterosteus aculeatus)* in Newfoundland and Labrador. *Journal of Fisheries Research Board*, Canada 27, 901–908.

Hogans, W. E., Dadswell, M. J., Uhazy, L. S., & Appy, R. G. (1993) Parasites of American shad, *Alosa sapidissima* (Osteichthyes: Clupeidae) from rivers of the North American Atlantic coast and the Bay of Fundy, Canada. Canadian Journal of Zoology 71, 941–946. 10.1139/z93-123.

Jabbar, A., Beveridge, I., & Bryant, M. S. (2015). Morphological and molecular observations on the status of *Crassicauda magna*, a parasite of the subcutaneous tissues of the pygmy sperm whale, with a re-evaluation of the systematic relationships of the genus *Crassicauda*. Parasitology Research 114, 835–841. 10.1007/s00436-014-4245-6.

Kalyaanamoorthy, S., Minh, B. Q., Wong, T. K. F., Von Haeseler, A., & Jermiin, L. S. (2017) ModelFinder: fast model selection for accurate phylogenetic estimate. Nature Methods 14, 587–589. 10.1038/nmeth.4285.

Kmentová, N., Hahn, C., Koblmüller, S., Zimmermann, H., Vorel, J., Artois, T., Gelnar, M., & Vanhove, M. P. M. (2021). Contrasting host–parasite population structure: morphology and mitogenomics of a parasitic flatworm on pelagic deepwater cichlid fishes from Lake Tanganyika. Biology 10, 797. 10.3390/biology10080797.

Klimpel, S., Seehagen, A., & Palm, H. W. (2003) Metazoan parasites and feeding behavior of four small-sized fish species from the central North Sea. Parasitology Research 91, 290–297. 10.1007/s00436-003-0957-8.

Ko, R. (1985) *Ascarophis minuta* n. sp. (Nematoda: Cystidicolidae) from a scorpaenid fish, *Sebasticus marmoratus*, in Hong Kong, southern China. Tropical Biomedicine 2, 99–105. 10.1645/GE-1169R1.1.

Ko, R. (1986) A preliminary review of Ascarophis (Nematoda) of fishes. Department of Zoology, University of Hong Kong, Occasional Publications 54 pp.

Køie, M. (2012) Nematode parasites in teleosts from 0 to 1540 m depth off the Faroe Islands (The North Atlantic). Ophilia 38, 217–243. 10.1080/00785326.1993.10429897.

Køie, M., Steffensen, J. F., Møller, P. R., & Christansen, J. S. (2008) The parasite fauna of *Arctogadus glacialis* (Peters) (Gadidae) from western and eastern Greenland. Polar Biology 31, 1010–1021. 10.1007/s00300-008-0440-1.

Kirjušina, M., & Vismanis, K. (2007) Checklist of the parasites of fishes of Latvia. FAO Fisheries Technical Paper 369/3, 106 pp.

Linstow, O. (1900) Die Nematoden. In: Fauna arctica: eine Zusammenstellung der arktischen Tierformen mit besonderer Beru cksichtigung des Spitzbergen-Gebietes auf Grund der Ergebnisse der Deutschen Expedition in das Nördliche Eismeer im Jahre 1898 4, 117–132.

Maggenti, A. (1981) General Nematology. Springer-Verlag, New York. 372 pp.

Meldal, B. H. M, Debenham, N. J, De Ley, P., De Ley, I. T., Vanfleteren, J. R., Vierstraete, A. R., Bert, W., Borgonie, G., Moens, T., Tyler, P. A., Austen, M. C., Blaxter, M. L., Rogers, A. D., & Lambsheae, P. J. D. (2007) An improved molecular phylogeny of the Nematoda with special emphasis on marine taxa. Molecular Phylogenetics and Evolution 42, 622–636. 10.1016/j.ympev.2006.08.025.

Miller, M. A., Pfeiffer, W., & Schwartz, T. (2010). Creating the CIPRES Science Gateway for inference of large phylogenetic trees. In 2010 Gateway Computing Environments Workshop (GCE), pp. 1–8. IEEE. 10.1109/GCE.2010.5676129.

Miscampbell, A. E., Lankester, M.W., & Adamson, M. L. (2004) Molecular and morphological variation within swim bladder nematodes, Cystidicola spp. Canadian Journal of Fisheries and Aquatic Science 61, 1143–1152. 10.1139/f04-064.

Moravec, F. (2007) Some aspects of the taxonomy and biology of adult spirurine nematodes parasitic in fishes: a review. Folia Parasitologica 54, 239–257. 10.14411/fp.2007.033.

Moravec, F. (2024) New data on the morphology of *Ascarophis parupenei* (Nematoda: Cystidicolidae), and intestinal parasite of the marine fish *Parupeneus indicus* in the Indian Ocean, revealed by SEM. Systematic Parasitology 101, 60. 10.1007/s11230-024-10188-7.

Moravec, F., Dykman, L. N., & Davis, D. B. (2024) Three new species of *Ascarophis* Van Beneden, 1871 (Nematoda: Cystidcolidae) from deep–sea hydrothermal vent fishes of the Pacific Ocean. Systematic Parasitology 101, 2. doi: 10.1007/s11230-023-10130-3.

Moravec, F., & González-Solís, D. (2007) Structure of the cephalic end of *Ascarophis mexicana* (Nematoda:Cystidicolidae), as revealed by SEM. Folia Parasitologica 54, 155–156.

Moravec, F., Hanzelová, V., & Gerdeaux, D. (2007) New data on the morphology of *Comephoronema oschmarini* (Nematoda, Cystidicolidae), a little-known gastrointestinal parasite of *Lota lota* (Teleostei) in Palaerarctic Eurasia. Acta Parasitologica 52, 135–141. 10.2478/s11686-007-0018-z.

Moravec, F., & Justine, J. L. (2007) A new species of *Ascarophis* (Nmeatoda, Cystidicolidae) from the stomach of the marine scorpaeniform fish *Hoplichthys citrinus* from a seamount off the Chesterfield Islands, New Caledonia. Acta Parasitologica 52, 238–246. 10.2478/s11686-007-0026-z.

Moravec, F., & Justine, J-L. (2009) Two cystidicolids (Nematoda, Cystidicolidae) from marine fishes off New Caledonia. Acta Parasitologica 54, 341–349. 10.2478/s11686-009-0058-7.

Moravec, F., & Justine, J-L. (2010) Two new genera and species of cystidicolids (Nematoda, Cystidicolida) from marine fishes off New Caledonia. Parasitology International 59, 198–205. 10.1016/j.parint.2010.01.005.

Moravec, F., & Justine, J. L. (2020) New records of spirurid nematodes (Nematoda, Spirurida, Guyanemidae, Philometridae and Cystidicolidae) from marine fishes off New Caledonia, with redescriptions of two species and erection of *Ichthyofilaroides* n. gen. Parasite 27, 5. 10.1051/parasite/2020003.

Moravec, F., & Nagasawa, F. (2018) *Rhabdochona angusticaudata* sp. n. (Nematoda: Rhabdochonidae) from the Japanese eel *Anguilla japonica*, and new records of some other nematodes from inland fishes in Japan. Folia Parasitologica 645, 016. 10.14411/fp.2018.016.

Moravec, F., Shamsi, S., & Justine, J-L. (2021) Redescription of *Ascarophis distorta* Fusco et Overstreet, 1978 (Nemadoda, Cystidicolidae) from the stomach of some butterflyfishes off New Caledonia. Acta Parasitologica 66, 907–914. 10.1007/s11686-021-00359-7.

Nadler, S. A., D’Amelio, S., Fagerholm, H. P., Berland, B., & Paggi, L. (2000) Phylogenetic relationships among species of *Contracaecum* Railliet & Henry, 1912 and *Phocascaris* Høst, 1932 (Nematoda: Ascaridoidea) based on nuclear rDNA sequence data. Parasitology 121, 455–463. 10.1017/S0031182099006423.

Nguyen, L-T., Schmidt, H. A., von Haeseler, A., & Minh, B. Q. (2014) IQ-TREE: a fast and effective stochastic algorithm for estimating maximum-likelihood phylogenies. Molecular Biology and Evolution 32, 268–274. 10.1093/molbev/msu300.

Nicoll, W. (1907) A contribution towards a knowledge of the Entozoa of British marine fishes. Pt. 1. Annals and Magazine of Natural History 19, 66–94.

Perdiguero-Alonso, D., Montero, F. E., Raga, J. A., & Kostadinova, A. (2008) Composition and structure of the parasite faunas of cod, *Gadus morhua* L. (Teleostei: Gadidae) in the North East Atlantic. Parasites & Vectors 1, 23. 10.1.1186/1756-3305-1-23

Pereira, A. N., Timi, J. T., Vierira, F. M., & Luque, J. L. (2012) A new species of *Neoascarophis* (Nematoda: Cystidicolidae) parasitic in *Mullus argentinae* (Perciformes: Mullidae) from the Atlantic coast of South America. Folia Parasitologica 59, 64–70. 10.14411/fp.2012.010.

Pereira, F. B., Pereira, A. N., & Luque, J. L. (2014) A new species of *Comephoronema* (Nematoda: Cystidicolidae) from the squirrelfish *Holocentrus adscensionis* (Beryciformes: Holocentridae) off Brazil Folia Parasitologica 61, 55–62. 10.14411/fp.2014.001

Petter, A. J. (1970) Quelque spirurides de poisons de la region Nantaise. Annales de Parasitologie 45, 31–46.

Polyanskii, Y. L. (1952) Some new and little known parasitic nematodes of the intestine of marine fishes. Trudy Zoologieschesko Instituta Akademiya Nauk SSSR 12, 133–147.

Polyanskii, Y. L. (1966) Parasites of the Fish of the Barents Sea. Trudy Zoologicheskogo Instituta. Akademii Nauk SSSR 19, 158 pp. Translated from Russian. Israel Program for Scientific Translations. Jerusalem.

Power, H., & Sedgwick, L. W. (1880) The new Sydenham Society’s Lexicon of Medicine and the allied sciences (Based on Mayne’s Lexicon). London 1879–1899. Vol. I, Part 4. [not paginated]

Prosser, S. W. J., Velarde-Aguilar, M. G., León-Règagnon, V., & Hebert, P. D. N. (2013) Advancing nematode barcoding: a primer cocktail for the cytochrome c oxidase subunit I gene from vertebrate parasitic nematodes. Molecular Ecology Resources 13.*6*, 1108–1115.

Punt, A. (1947) Quelques nematodes parasites de poisons de la mer du nort. II. *Bulletin du Musé de Rouyale d’Histoire Naturelle*, Belge 23, 1–13.

Rahman, H. (1966) *Ascarophis crassicollis* Dollfus and Campana–Rouget in Scottish waters. Annals and Magazine of Natural History, Year 1965, Ser. XIII, 8 (87-88), 187–192.

Rambaut, A., Drummond, A. J., Dixie, X., Bule, G., & Scuhard, M. H. (2018) Posterior summarization in Bayesian phylogenetics using Tracer 1.7. Systematic Biology 67, 901–904.

Rasheed, S. (1965) Observations on the spiuroid nematodes of fish with a revision of the genus *Metabronema* Yorke and Maplestone, 1926. Zeischrift für Zoologie Systematik Evolutionsforsch 3, 359–387.

Rees, G. (1945) A record of parasitic worms from fishes in rock pools at Aberystwyth. Parasitology 36, 165–167.

Reubens, J., Degraer, S., & Vinex, M. (2010) Spatial and temporal movements of cod (*Gadus morhua*) in a wind farm in the Belgium part of the North Sea using acoustic telemetry, a VPS study. Offshore Wind Farms in The Belgium Part of the North Sea 3, 39–476.

Rodrigues, H., Varela, M. C., Rodrigues, S. S. & Cristófaro, R. (1973) Alguns nematódeos de peixes do oceano Atlântico - cost continental Portugue sa e costa do Norte da África. Memorias Instituto Oswaldo Cruz 71, 247–256.

Ronquist, F., Teslenko, M., Van der Mark, P., Ayres, D. L., Darling, A., Hohna, S., Larget, B., Liu, L., Suchard, M. A., & Huelsenbeck, J. P. (2012). MrBayes 3.2: Efficient Bayesian phylogenetic inference and model choice across a large model space. Systematic Biology 61, 539–542. 10.1093/sysbio/sys029.

Rossin, M. A., Incorvaia, I. A., & Timi, J. T. (2012) A new species of *Neoascarophis* (Nematoda: Cystidicolidae) parasitic in *Macrourus carinatus* (Macrouridae) from Argentinean waters. Journal of Parasitology 98, 643–647. 10.1645/JP-GE-2947.1

Saiki, R. K., Gelfand, D. H., Stoffel, S., Scharf, S. J., Higuchi, R., Horn, G. T, Mullis, K. B., & Erlich, H. A. (1988). Primer–directed enzymatic amplification of DNA with a thermostable DNA polymerase. Science 239, 487–491. 10.1126/SCIENCE.2448875,

Scott, J. S. (1981) Alimentary tract parasites of haddock (*Melanogrammus aeglefinus* L.) on the Scotian Shelf. Canadian Journal of Zoology 59, 2244–2252.

Skrjabin, K. I., Sobolev, A. A., & Ivashkin, V. M. (1967) [Essentials of Nematodology, Volume XVI Spirurata of animals and man and the diseases caused by them Part 4. Thelazioidea (edited by Skrjabin KI) Academy of Sciences of the USSR, Moscow, 624 pp. (Israel Program of Scientific Translation Ltd. 1971).

Sobecka, E., & Łuczak, E. (2015) Species richness and the diversity of parasite communities of eelpout *Zoarces viviparus* (Linnaeus, 1758) in the Oder River estuary, Poland. Oceanological and Hydrobiological Studies 44, 520–529.

Sokolov, S. G., Voropaeva, E. L., & Malysheva, S. V. (2019) Redescription and molecular characterization of *Comephoronema werestschagini* Layman, 1933 (Nematoda: Cystidicolidae) from the endemic Baikal fish *Cottocomephorus grwingkii* (Dybowski, 1874) (Scorpaeniformes: Cottocomephoridae) with some comments on cystidicolid phylogeny. Russian Journal of Nematology 27, 57–66. 10.24411/0869-6918-2019-10007.

Srivastava, L. P. (1966) The helminth parasites of the five-bearded rockling, *Onos mustelus* (L.) from the shore at Mumbles Head, Swansea. Annals and Magazine of Natural History 9, 469–480.

Studnicka, M. (1965) Internal parasites of the cod, *Gadus callarias* L., from the Gdansk Bay of the Baltic Sea. Acta Parasitologica Polonica 13, 283–290.

Van Beneden, P-J. (1870) Les poisons des côtes de Belgique, leurs parasites et leurs commensaux. Hayez Bruxelles 38, 100 pp., 9 Plates.

Vidal, V., Ortiz, J., Diaz, J. I, Zafrilla, B., Bonete, M. J, De Ybañez, M. R., & Barbosa, A. (2016). Morphological, molecular and phylogenetic analyses of the spirurid nematode *Stegophorus macronectes* (Johnston & Mawson, 1942). Journal of Helminthology 90, 214–222. 10.1017/S0022149X15000218.

Wong, W. L., Tan, W. B., & Lim, H. S. (2006) Sodium dodecyl sulfate as a rapid clearing agent for studying the hard parts of monogeneans and nematodes. Journal of Helminthology 80, 87–90. 10.1079/joh2005320

Yorke, W., & Maplestone, P. A. (1926) *The nematode parasites of vertebrates*. P. Blakiston Zhukov, E. V. (1960) Endoparasitic worms of fishes from the Sea of Japan and South Kurile Shoal. Trudy Zoologicheskogo Instituta. Akademii Nauk SSSR 28, 3–146. (In Russian with English summary).

Zubchenko, A. V. (1980) Parasitic fauna of Anarhichadidae and Pleuronectidae families of fish in the northwest Atlantic. *International Commission for Northwest Atlantic Fisheries*, Selected Papers 6, 41–46.

Zubchenko, A. V. (1987) Vertical zones and formation of the parasitic fauna in deepwater fishes from off-shore areas of the North Atlantic. *Northwest Atlantic Fisheries Organization, Scientific Council Report* Document 84/41, N1326, June. 22 pp.

Zubchenko, A. V., & Karasev, A. B. (1986) Parasitofauna of Barents Sea marine fish. In Nefedov, N. F. Conditions concerning ichthyofauna in the Barents Sea. Murmansk Marine Biological Institue, Apatity Akademia Nauk SSSR pp. 132–151.

